# Biophysical models reveal the relative importance of transporter proteins and impermeant anions in chloride homeostasis

**DOI:** 10.1101/216150

**Authors:** Kira M. Düsterwald, Christopher B. Currin, Richard J. Burman, Colin J. Akerman, Alan R. Kay, Joseph V. Raimondo

## Abstract

Fast synaptic inhibition in the nervous system depends on the transmembrane flux of Cl^−^ ions based on the neuronal Cl^−^ driving force. Established theories regarding the determinants of Cl^−^ driving force have recently been questioned. Here we present biophysical models of Cl^−^ homeostasis using the pump-leak model. Using numerical and novel analytic solutions, we demonstrate that the Na^+^/K^+^-ATPase, ion conductances, impermeant anions, electrodiffusion, water fluxes and cation-chloride cotransporters (CCCs) play roles in setting the Cl^−^ driving force. Our models, together with experimental validation, show that while impermeant anions can contribute to setting [Cl^−^]_i_ in neurons, they have a negligible effect on the driving force for Cl^−^ locally and cell-wide. In contrast, we demonstrate that CCCs are well-suited for modulating Cl^−^ driving force and hence inhibitory signalling in neurons. Our findings reconcile recent experimental findings and provide a framework for understanding the interplay of different chloride regulatory processes in neurons.

## Introduction

Fast synaptic inhibition in the nervous system is mediated by type A γ-aminobutyric acid receptors (GABA_A_Rs) and glycine receptors (GlyRs), which are primarily permeable to chloride (Cl^−^) (Farrant and Kaila, 2007). Together with the neuronal membrane potential, the transmembrane gradient for Cl^−^ sets the driving force for Cl^−^ flux across these receptors, controlling the properties of inhibitory signaling. Modification of neuronal intracellular Cl^−^ concentration ([Cl^−^]_i_) has been shown to play a causative role in multiple neurological diseases including epilepsy, chronic pain, schizophrenia and autism (Rivera et al., 2004; Huberfeld et al., 2007; Price et al., 2009; Hyde et al., 2011; Tyzio et al., 2014). Similarly, intracellular Cl^−^ is thought to be modulated during brain development so that GABAergic transmission contributes optimally to the construction of neural circuits (Ben-Ari, 2002). Given the importance of Cl^−^ for brain function and dysfunction, the cellular mechanisms that control its transmembrane gradient and driving force are of considerable interest.

Plasmalemmal Cl^−^ transporters, in particular cation-chloride cotransporters (CCCs), are understood to be the major mechanism by which neurons regulate the driving force for Cl^−^ permeable anion channels (Kaila et al., 2014). Recently, it has been suggested that in fact impermeant anions control local [Cl^−^]_i_ and driving force (Glykys et al., 2014), rather than the CCCs. The majority of intracellular anions are impermeant to the neuronal membrane; these include ribo- and deoxynucleotides, intracellular proteins and metabolites (Burton, 1983). Impermeant anions induce what is known as the Donnan (or Gibbs-Donnan) effect (Hill, 1956; Sperelakis, 2012) – an uneven distribution of impermeant molecules across the membrane which is osmotically unstable. Without active ion transport to counter this effect, neurons would swell and burst (Kay, 2017). Animal cells, including neurons, maintain cell volume in the presence of impermeant anions by using the Na^+^/K^+^-ATPase to pump Na^+^ out of the cell and K^+^ in, along with the passive movement of water and other ions (Tosteson and Hoffman, 1960; Armstrong, 2003; Liang et al., 2007; Kay, 2017). This pump-leak mechanism, whilst stabilizing cell volume, also establishes the negative resting membrane potential and transmembrane Na^+^ and K^+^ gradients, which serve as energy sources for the coupled transport of other molecules, including Cl^−^ by CCCs. In the absence of active Cl^−^ transport, [Cl^−^]_i_ is set by the membrane potential; i.e. the Nernst potential of Cl^−^ 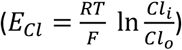 equals the transmembrane potential (V_m_).

The transmembrane Cl^−^ gradient and the driving force (DF=V_m_-E_Cl_) for Cl^−^ permeable ion channels are therefore the outcome of multiple, dynamically interacting mechanisms. This makes experimental investigation of the determinants of Cl^−^ driving force difficult, particularly at a local level. Computational models based on established biophysical first principles are productive means for exploring the roles of cellular mechanisms in generating local Cl^−^ driving force. Here, we establish numerical and novel analytic solutions for an inclusive model of Cl^−^ homeostasis to elucidate the determinants of the neuronal driving force for Cl^−^.

We demonstrate that baseline [Cl^−^]_i_ is a product of the interaction of Na^+^/K^+^-ATPase activity, the average charge of impermeant anions, ion conductances, CCCs and water permeability. Consistent with recent experimental reports (Glykys et al., 2014), and our own experimental validation using electroporation of anionic dextrans and optogenetic probing of E_GABA_, we find that impermeant anions can contribute to setting [Cl^−^]_i_. However, we find that they can only affect the Cl^−^ driving force by modifying active transport mechanisms, and then only negligibly. Impermeant anions therefore do not appreciably modify synaptic signaling properties, contrary to recent experimental interpretation (Glykys et al., 2014). In contrast, we demonstrate using biophysical models and gramicidin perforated-patch clamp recordings that CCCs selectively regulate substantial changes in the Cl^−^ driving force. This is consistent with a meta-analysis of experimental data from the field, which shows a strong correlation between the activity of the specific CCC, KCC2, and Cl^−^ driving force. The ability of CCCs to specifically modulate Cl^−^ at a local level depends on the characteristics of Cl^−^ electrodiffusion in the structure concerned, demonstrated using multicompartment modelling. Together, our models provide a theoretical framework for understanding the interplay of chloride regulatory processes in neurons and interpreting experimental findings.

## Results

### A biophysical model based on the pump-leak mechanism demonstrates the importance of the sodium-potassium ATPase for setting transmembrane ion gradients including chloride

To compare the effects of impermeant anions and Cl^−^ cotransport on Cl^−^ homeostasis we first developed a single compartment model based on the pump-leak formulation (Tosteson and Hoffman, 1960; Kay, 2017) (Fig. 1A). This model, defined by a set of differential equations, incorporated mathematical representations of the three major permeable ion species Cl^−^, K^+^, Na^+^ as well as impermeant anions (X^z^) with average charge z. Permeable ions could move across the cellular membrane via passive conductances according to each ion’s respective electrochemical gradient. Further, active transport of Na^+^ and K^+^ by the Na^+^/K^+^-ATPase and cotransport of Cl^−^ and K^+^ by the cation-chloride cotransporter KCC2 with a non-zero conductance g_KCC2_ of 20 µS/cm^2^ unless otherwise stated were included. Finally, our formulation accounted for the dynamics of cell volume (w), intracellular osmolarity (Π) and transmembrane voltage (V_m_) (see Materials and Methods). Importantly, regardless of initial starting concentrations of permeant or impermeant ions, cell volume or V_m_, the model converged to stable fixed points without needing to include any means for ‘sensing’ ion concentration, volume or voltage. In the case of permeable ions, initial concentrations did not even determine the final cellular volume. For example, despite initiating the model with different starting concentrations of Cl^−^ (1, 15, 40 and 60 mM respectively, Fig. 1B), the [Cl^−^]_i_ always converged to the same stable concentration of 5.2 mM, a typical baseline [Cl^−^]_i_ for adult neurons, and volume always converged to 2.0 pL, a typical volume for hippocampal neurons (Ambros-Ingerson and Holmes, 2005). The model is robust in the sense that its convergence to a stable steady state does not depend on a narrow set of parameters and initial values. Another proposed model of the Na^+^/K^+^-ATPase with experimental validation (Hamada et al., 2003) parameterized for our steady state values had similar activity in terms of net flux rate in our range of [Na^+^]i and invariance to initial permeable ionic concentrations, (**Figure 1 – figure supplement 1A-B**).

**Figure 1:**
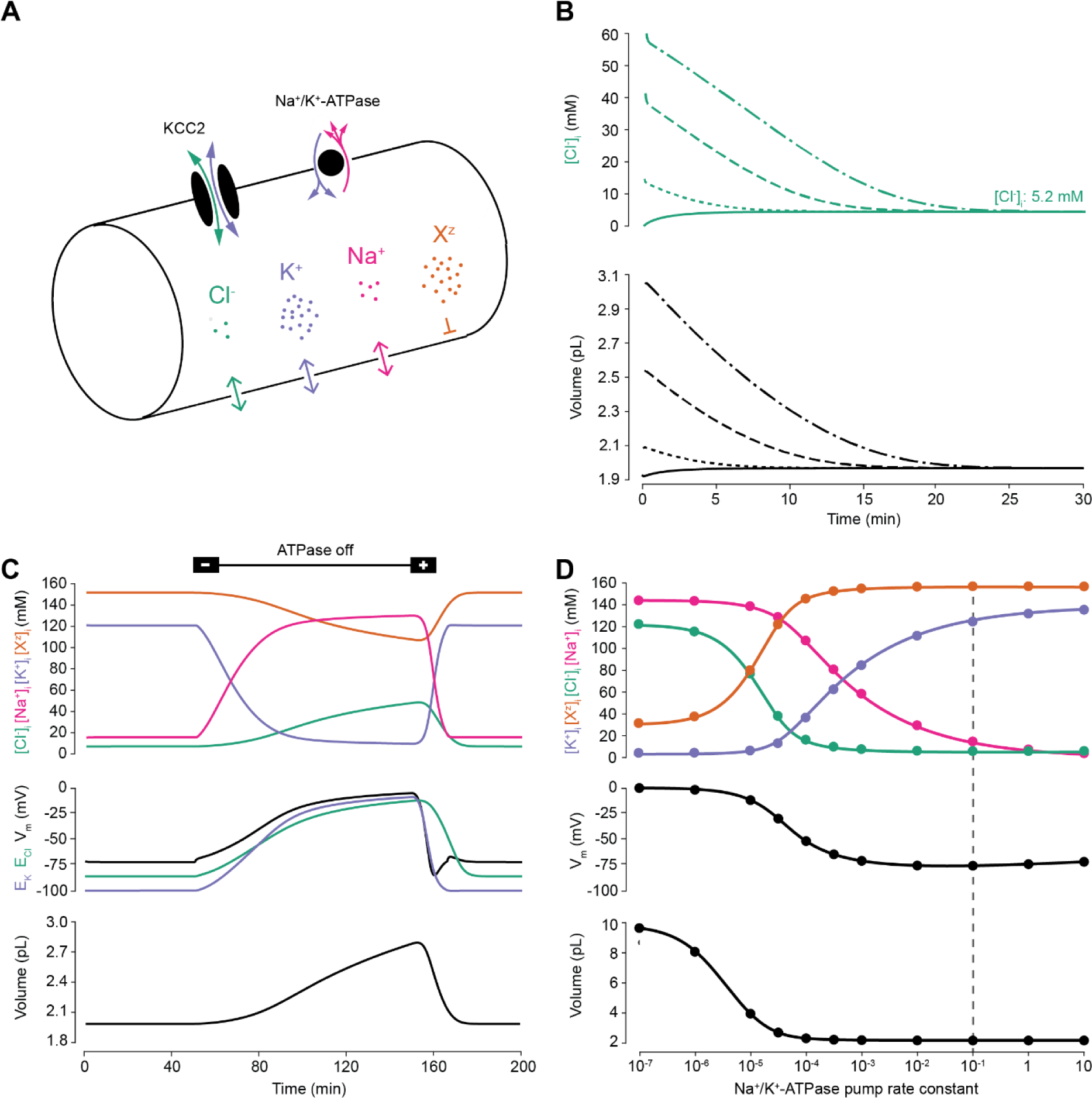
A biophysical model of ion dynamics based on the pump-leak mechanism demonstrates the importance of the sodium-potassium ATPase for setting transmembrane ion gradients including chloride. *(A) A single cell compartment was modelled as a cylinder with volume changes equivalent to changes in cylindrical radius. Dynamics of membrane permeable potassium (purple, K^+^), sodium (pink, Na^+^) and chloride (green, Cl^−^) ions were included. Impermeant anions (orange, X^z^) had an average intracellular charge z. The KCC2 transporter moved Cl^−^ and K^+^ in equal parts according to the transmembrane gradient for the two ions. The sodium-potassium ATPase transported 3 Na^+^ ions out for 2 K^+^ ions moved into the cell. (B) Regardless of intracellular starting concentrations of the permeable ions, the model converged to identical steady state values for all parameters without needing to include any means for ‘sensing’ ion concentration, volume or voltage. We show the result for Cl^−^ as a time series of [Cl^−^]_i_ (top panel) and volume (bottom panel). (C) The ATPase plays a key role in maintaining steady state ion concentration, membrane voltage (V_m_) and volume. Switching off the ATPase results in a continuous increase in cell volume (bottom panel), membrane depolarization (middle panel) and ion concentration dysregulation (top panel with colours per ion as in ‘A’. All cellular parameters recovered when the ATPase was reactivated. (D) The model’s analytic solution showed exact correspondence with steady state values generated by numerical, time series runs (dots) for varying ATPase pump rates. Steady state values for the concentrations of the ions with colours as in A (top panel); V_m_ (middle panel) and volume (bottom panel). The dashed line indicates the default ATPase pump rate used in all simulations unless specified otherwise. The result in (B) is replicated with Hamada et al.’s experimentally validated model of the ATPase in Figure 1 – figure supplement 1B, and the models’ respective pump fluxes are compared in figure supplement 1A*.

Consistent with previous results (Xiao et al., 2002; Dierkes et al., 2006; Dijkstra et al., 2016), ‘turning off’ the activity of the Na^+^/K^+^-ATPase in our model led to a progressive collapse of transmembrane ion gradients, progressive membrane depolarization and continuous and unstable cell swelling. Such effects could be reversed by reactivation of the pump (Fig. 1C). The relative activity of the Na^+^/K^+^- ATPase sets the final stable values for ion concentrations (of both permeable and impermeant ions), V_m_ and volume (Fig. 1C). When we increased the activity of the Na^+^/K^+^-ATPase in our model, the final steady-state concentration for K^+^ increased, whilst Na^+^ and Cl^−^ dropped to levels that approximate those observed in mature neurons (Fig. 1D). At the same time, the final, stable-state membrane potential and cell volume also decreased. Interestingly, as has been observed previously (Fraser and Huang, 2004), beyond a certain level, further increases in Na^+^/K^+^-ATPase activity have negligible effects on cell volume and transmembrane voltage. For subsequent analysis, we chose a ‘default’ effective pump rate for the Na^+^/K^+^-ATPase of approximately 1.0 × 10^−2^ C/(dm^2^.s), that is a pump rate constant of 10^−1^ C/(dm^2^.s) (equation (2)), and average intracellular impermeant anion charge (z) of −0.85 as extrapolated from reasonable cellular ionic concentrations and osmolarity (Lodish et al., 2009; Raimondo et al., 2015). This resulted in steady-state ion concentrations and membrane potentials that approximate those experimentally observed in mature neurons: Cl^−^ 5 mM; K^+^ 123 mM; Na^+^ 14 mM; X^z^ 155 mM; and a V_m_ of −72.6 mV (Jiang and Haddad, 1991; Diarra et al., 2001; Tyzio et al., 2008). We were able to corroborate the numerical solutions for final steady-state values by developing a parametric-analytic solution (**Supplementary file 1**). We observed exact correspondence between the numerical and analytic solutions within our model (Fig. 1D). In subsequent analyses, this novel analytic solution allowed us to explore rapidly a large parameter space to determine how various cellular attributes might affect Cl^−^ homeostasis.

### Membrane chloride conductance affects steady-state intracellular chloride concentration only in the presence of cation-chloride cotransport

Using the analytic solution, we investigated how changes in baseline ion conductance for the major ions in our model (g_K_, g_Na_ and g_Cl_) affected Cl^−^ homeostasis. We calculated the steady-state values for the Cl^−^ reversal potential (E_Cl_) and potassium reversal potential (E_K_), resting membrane potential (V_m_) and volume (w) whilst independently manipulating the conductance for each ion (Fig. 2). Increasing the baseline potassium conductance (g_K_) resulted in E_Cl_, E_K_ and V_m_ converging to similar steady-state values (Fig. 2A) without significantly affecting cell volume. We were also able to replicate the classic dependence of membrane potential on log([K^+^]_o_) (**Figure 2 – figure supplement 1**). In contrast, increasing the baseline sodium conductance (g_Na_) beyond 20 µS/cm^2^ resulted in a steady depolarization of E_Cl_, E_K_ and V_m_ with accompanying cell swelling (Fig. 2B).

**Figure 2:**
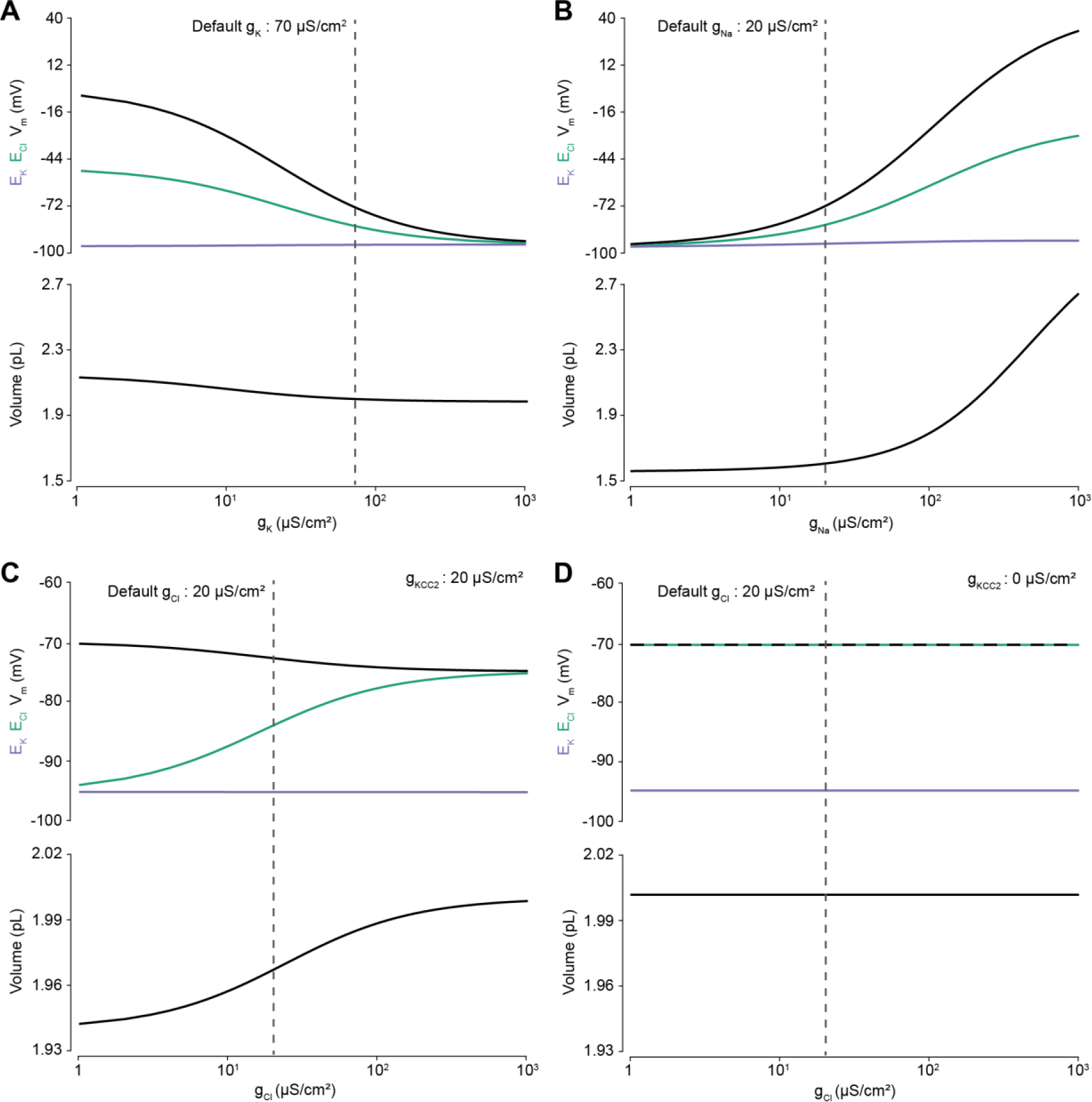
Membrane chloride conductance affects steady-state intracellular chloride concentration only in the presence of cation-chloride cotransport. *Steady state values for different ionic conductance were calculated using the model’s analytic solution. (A) Steady state E_Cl_ (green), E_K_ (purple), V_m_ (black) and volume w (bottom panel) were calculated at different K^+^ conductances (g_K_). Increasing gK resulted in a convergence of steady state E_Cl_, E_K_ and V_m_. (B) Increasing Na^+^ conductance g_Na_ resulted in a progressive increase in steady state E_Cl_, E_K_ and V_m_ with accompanying cell swelling. (C) In the presence of active cation-chloride cotransport (g_KCC2_ =* 20 µS/cm^2^), *increasing Cl^−^ conductance shifted steady state E_Cl_ from E_K_ toward V_m_. (D) In the absence of KCC2 activity, g_Cl_ had no effect on steady state parameters. E_Cl_ equals V_m_ in all instances. Dashed lines indicate the default values for g_K_, g_Na_ and g_Cl_. In Figure 2 – figure supplement 1 the classic dependence of the membrane potential on log([K^+^]_o_) is shown*.

The effect of manipulating Cl^−^ conductance (g_Cl_) depended on the activity of concurrent cation-chloride cotransport by KCC2 (Fig. 2C). In the presence of active KCC2 at very low values of g_Cl_, the steady state [Cl^−^]_i_ is such that E_Cl_ approaches E_K_. This follows because in the absence of alternative Cl^−^ fluxes, KCC2 utilizes the transmembrane potassium gradient to transport Cl^−^ until E_Cl_ equals E_K_. With increasing g_Cl_ however, E_Cl_ increases, moving away from E_K_ towards V_m_, and at very high Cl^−^ conductances E_Cl_ and V_m_ approached similar values in our model. Without the activity of KCC2, g_Cl_ had no effect on steady state E_Cl_, E_K_, V_m_ or volume, with gcl able to be very low but greater than zero (Fig. 2D). In this instance E_Cl_ always equals V_m_ as the movement of Cl^−^ across the membrane is purely passive. Without the activity of KCC2, there can be no driving force for Cl^−^ flux at steady state (V_m_-E_Cl_ = 0). Our model therefore behaved in a manner consistent with established theoretical predictions (Kaila et al., 2014).

### Cation-chloride cotransport sets the chloride reversal and driving force for transmembrane chloride flux

Next, we used our single-cell unified model to explore how the activity of cation-chloride cotransport affects Cl^−^ homeostasis. In our model, the activity of KCC2 is set by the conductance of KCC2 (g_KCC2_). Using the numerical formulation with the default values described in Figure 1 and 2, we steadily increased g_KCC2_ from 20 µS/cm^2^ to 370 µS/cm^2^ and tracked changes to E_Cl_, E_K_, V_m_ and volume. Increasing KCC2 activity over time caused a steady decrease in [Cl^−^]_i_ reflected by a hyperpolarization of E_Cl_ (Fig. 3A). V_m_ decreased only modestly, resulting in an increase in the driving force for Cl^−^ flux that tracks the increase in g_KCC2_. This effect saturates as E_K_ constitutes a lower bound on E_Cl_. Importantly, increases in g_KCC2_ resulted in persistent changes to E_Cl_ and the driving force for Cl^−^. Employing alternate models for KCC2 (Fraser and Huang, 2004; Lewin et al., 2012; Raimondo et al., 2012) and the ATPase (Hamada et al., 2003) did not change this result when compensation for parameterization was given, although different KCC2 models result in different kinetic rates for Cl^−^ and K^+^ transport (**Figure 3 – figure supplement 1 and 2**). Using the analytic solution to our model we calculated how KCC2 activity affects steady state values of E_Cl_, E_K_, V_m_, w and Cl^−^ driving force (Fig. 3B). In confirmation of our findings in Figure 2D, with no KCC2 activity (g_KCC2_ = 0), E_Cl_ equaled V_m_ and the Cl^−^ driving force was zero. As we increased g_KCC2_, steady state E_Cl_ pulled away from V_m_ and approached E_K_. This resulted in an increase in Cl^−^ driving force (V_m_-E_Cl_) with steady state values of 11.3 mV at our chosen default value of g_KCC2_. The results obtained with our model are therefore fully consistent with the view that CCCs, in this case KCC2, establish the driving force for Cl^−^.

**Figure 3:**
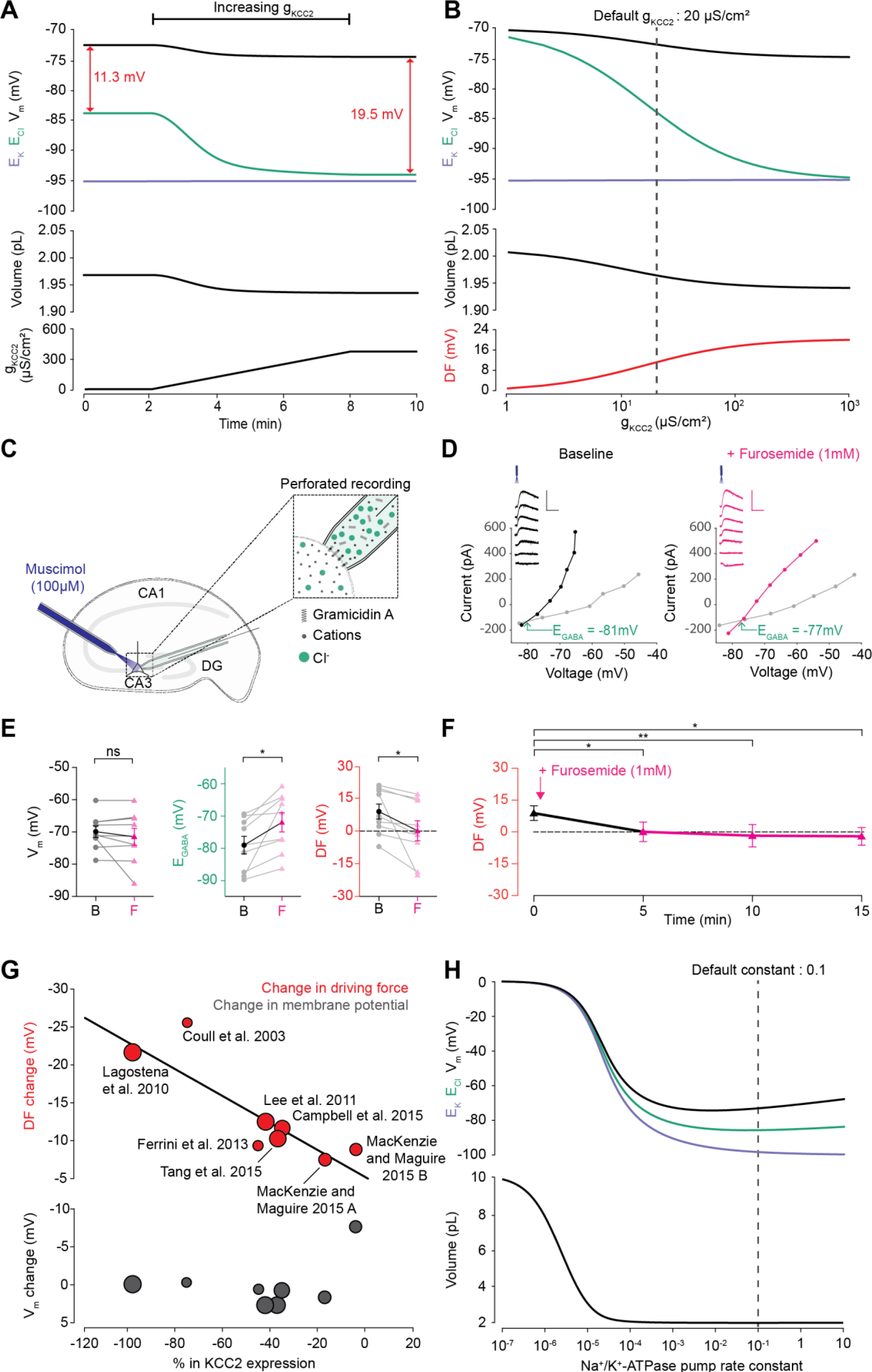
Cation-chloride cotransport sets the chloride reversal and driving force for transmembrane chloride flux. *(A) Increasing KCC2 activity in our model by increasing g_KCC2_ from 20 µS/cm^2^ to 370 µS /cm^2^ resulted in a persistent decrease in E_Cl_ (green), a minimal decrease in E_K_ (purple), V_m_ (black) and volume w (bottom panel). This resulted in a permanent increase in the DF for Cl^−^ via a change in E_Cl_ from −83.9 mV to −93.2 mV (red). (B) The steady state values for E_Cl_, E_K_, V_m_ (top panel), volume (middle panel) and Cl^−^ driving force (DF, bottom panel, red) at different KCC2 conductances. Increasing KCC2 activity resulted in a decrease in steady state E_Cl_ and DF. E_K_ represented a lower bound on E_Cl_ at high KCC2 conductances. Similar results were noted for other kinetic models of KCC2 (figure supplement 1 and 2). (C) Schematic showing experimental setup. Gramicidin perforated patch-clamp recordings were performed on CA3 pyramidal cells from rat hippocampal organotypic brain slices. (D) Insets depict GABA_A_R currents elicited by somatic application (20 ms) of muscimol (10 μM) at different voltages. Calibration: 500 ms, 500 pA. Current-voltage (IV) plots with holding current (reflecting membrane current, grey) and total current (reflecting membrane current plus the muscimol-evoked current, black or pink) were drawn to calculate changes in V_m_, E_GABA_ and DF before (left, black) and after (right, pink) furosemide was applied. Voltages were corrected for series resistance error. (E) Population data showing significant changes in E_GABA_ and DF but not V_m_ five minutes after furosemide application. (F) Changes in DF over time show a significant decrease from baseline once furosemide was introduced. (G) Meta-analysis of experimental studies demonstrates a correlation between KCC2 activity (% change) and Cl^−^ DF (mV, top plot, red) but not membrane potential (mV, bottom plot, grey), confirming the role of KCC2 for establishing the neuronal Cl^−^ gradient in adult tissue. The data and scoring system used to generate the regression can be found in Supplementary file 2 (Tables S2-1 and S2-2). (H) Sufficient Na^+^/K^+^-ATPase activity is critical for steady state ionic gradients, V_m_ (top panel, colours as in ‘A’) and volume (bottom panel), but these variables are relatively stable near the default pump rate (dashed line). ‘ns’, non-significant; *p <0.05; ** p < 0.01*.

To test this theoretical finding, we performed gramicidin perforated patch-clamp recordings from CA3 hippocampal neurons in rat organotypic brain slices whilst activating Cl^−^ permeable GABA_A_ receptors with muscimol (10 µM), in order to measure the GABA_A_R driving force, which approximates Cl^−^ driving force (Fig. 3C). We then tested our model predications by applying the CCC blocker, furosemide (1 mM) (Fig. 3D, E). We noted that after furosemide was introduced the E_GABA_ became significantly more depolarized (baseline: −78.81 ± 2.78 mV vs furosemide: −71.59 ± 3.04 mV, *n* = 10, *p* = 0.01, *paired t-test*) whilst there was no significant difference in V_m_ (−69.92 ± 1.77 mV vs −71.51 ± 2.61 mV, *n* = 10, *p* = 0.36, *paired t-test*) (Fig. 3E). This reflects a significant change in the GABA_A_R driving force (8.89 ± 3.44 mV vs 0.08 ± 4.57 mV, *n* = 10, *p* = 0.04, *paired t-test*). Furthermore, we then noted that this change in driving force persisted for at least 15 minutes post application of furosemide (Fig. 3F). These results are consistent with our model predictions and demonstrate how the application of a CCC blocker reduces the GABA_A_R driving force (and hence Cl^−^ driving force) by selectively depolarizing E_GABA_ with negligible effects on V_m_.

In addition, we sought experimental data from the literature to determine whether changes in KCC2 activity correlate with alterations to steady-state [Cl^−^]_i_. We focused on changes in KCC2 expression level, as this is likely to be a strong predictor of changes in KCC2 activity. Indeed, in a meta-analysis of 7 studies and 8 experiments from our review of 26 studies, weighted for methodological biases and data quality, we observed a significant correlation (R^2^ = 0.796, p < 0.001) between the change in KCC2 expression and Cl^−^ driving force (Fig. 3G). Absolute changes in V_m_ were less than 2 mV in all but one study, meaning that the change in driving force could be ascribed to significant shifts in E_GABA_ (R^2^ = 0.045, p < 0.001). The outlier data point (showing a 8.45 mV change) was from a study into the effects of acute stress, where other factors could have transiently influenced V_m_ (MacKenzie and Maguire, 2015). The meta-analysis supports the prediction that cation-chloride cotransport by KCC2 is an important determinant of [Cl^−^]_i_ (R^2^ = 0.83, p < 0.001, 9 studies) and driving force (see **Supplementary file 2 Table S2-1** for raw data, and the scoring table for weighting in **Table S2-2**).

Since Cl^−^ cotransport by KCC2 depends on the transmembrane gradient for K^+^, which in turn is established by the Na^+^/K^+^-ATPase, we explored how the Na^+^/K^+^-ATPase pump rate affected E_Cl_ in the presence of KCC2 activity (g_KCC2_ = 20 µS/cm^2^). As depicted in Fig. 3H, the Na^+^/K^+^-ATPase pump is a critical determinant of intracellular Cl^−^ homeostasis. Of interest, above a pump rate constant of 10^−3^ C/(dm^2^.s), where cell volume is stable, even many fold changes in pump rate have minimal effects on E_Cl_ and V_m_. It therefore seems unlikely that neurons would adjust the Na^+^/K^+^-ATPase to modulate Cl^−^ driving force.

### Altering the concentration of intracellular or extracellular impermeant anions, without changing the average charge of impermeant anions, does not affect the steady state gradient or driving force for chloride

To determine the effect of impermeant anions on Cl^−^ homeostasis we first explored whether adjusting the concentration of impermeant anions ([X]_i_), while maintaining a constant average impermeant anion charge (z), had any impact on E_Cl_, E_K_, V_m_ or volume. We initiated the full single-compartment model with different starting concentrations of impermeant anions (but the same average charge, z = −0.85), and observed that regardless of the initial concentration of impermeant anions, over a period of minutes, the cell adjusted its volume to give an identical steady-state impermeant anion concentration (Fig. 4A, [A]_i_ = 155 mM). Analytically, it can be shown that the number of moles of X determines completely the volume of the compartment, while the permeant ions alone cannot be used to predict steady state volume (Kay, 2017). Similarly, all initial impermeant anion concentrations resulted in identical steady state values of E_Cl_ (−83.8 mV), E_K_ (−95.1 mV) and V_m_ (−72.6 mV) (Fig. 4B). This shows that simply adjusting the amount of impermeant anions within a cell has no persistent effect on [Cl^−^]_i_.

**Figure 4:**
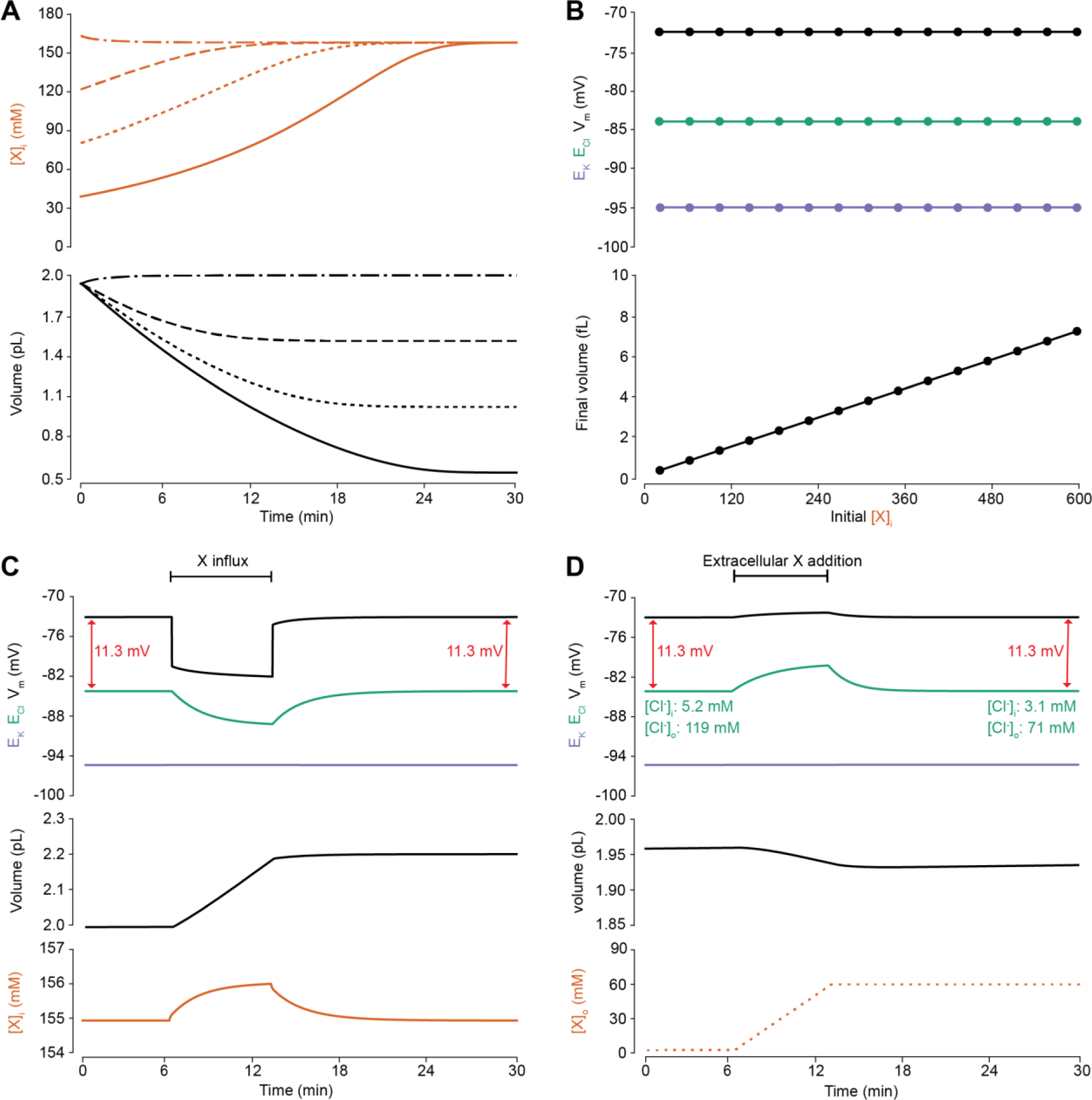
Adding intracellular or extracellular impermeant anions, without changing the average charge of impermeant anions, does not affect the steady state gradient or driving force for chloride. *(A) Initiating the model with different starting concentrations of intracellular impermeant anions ([X]_i_) with the same average charge z (orange, top panel), led to compensatory volume changes (bottom panel) which resulted in identical steady state concentrations. (B) Steady state E_Cl_ (green), E_K_ (purple) and V_m_ (black) were identical regardless of initial [X]_i_. Final volume however was a linear function of initial [X]_i_ (bottom panel). (C) Addition of impermeant anions of the same average charge (z) caused transients shifts in E_Cl_ (green, top panel), E_K_ (purple), V_m_ (black) as well as [X]_i_ (orange, bottom panel) for the duration of the influx, and sustained increases in volume (black, middle panel). No persistent changes in E_Cl_, E_K_ or V_m_ were observed. The full time-dependent ionic and water fluxes for the experiment are shown in Figure 4 – figure supplement 1, which shows that the inward flux of impermeant anions causes fluxes of all other ions. (D) Similarly, the addition of extracellular impermeant anions in an osmoneutral manner causes transient shifts in the permeable ion gradients (top panel, colours as in ‘C’), and sustained changes in cellular volume (black, middle panel) as well as the extracellular X and extracellular and intracellular Cl^−^ concentrations.*

We then tested the effect of dynamically adding impermeant anions with the default average charge either intracellularly (Fig. 4C) or extracellularly (Fig. 4D). While impermeant anions are being added to the cell, the membrane potential hyperpolarizes and E_Cl_ decreases. However, following the cessation of impermeant anion influx, E_Cl_, E_K_, V_m_ and [X]_i_ return to steady state values due to compensatory changes to cell volume (Fig. 4C). There are transient transmembrane fluxes of all ions while anions are added into the cell, and in particular the inward flux of the cations Na^+^ and K^+^, such that the sum [X]_i_ + [Cl^−^]_i_ is not necessarily kept constant during impermeant anion addition (**Figure 4 – figure supplement 1**). Impermeant ions (with default average charge) were added to the extracellular space, which is effectively an infinite bath in the model, while proportional decreases in [Cl^−^]_o_ were applied to correct for the changes to charge and osmotic balance. Additions in the extracellular space, similarly, resulted in a temporary depolarization of E_Cl_ and V_m_, but no persistent shift in these parameters (Fig. 4D). The addition of extracellular impermeant anions did however result in a small compensatory decrease in cell volume secondary to the large shifts in [Cl^−^]_i_ required to maintain the proportion of [Cl^−^]_i_ to [Cl^−^]o according to the Nernst potential. In summary, there is no lasting effect on the reversal potential or driving force for Cl^−^ if only the concentration of a neuron’s intracellular or extracellular impermeant anions is altered.

### Changing the average charge of impermeant anions can drive substantial shifts in the reversal potential for chloride, but has negligible effects on chloride driving force

We next sought to determine how changes in the average charge of the impermeant ions (z) might influence the driving force on Cl^−^. Such changes in average z could be associated with various cellular processes, including post-translational modifications of proteins that decrease their charge without changing the absolute number of protein molecules. To investigate this parameter, we modified the average z of intracellular impermeant anions from −0.85 to −1 whilst measuring accompanying changes in E_Cl_, E_K_, V_m_ and cell (Fig. 5A). We found that this shift to a more negative average z resulted in both a transient and persistent decrease in E_Cl_, E_K_ and V. Importantly, the shifts in E_Cl_ were accompanied by broadly matching shifts in E_K_ and V, which resulted in a small change in the driving force for Cl^−^ of < 0.2 mV. Both numerical and analytic calculation of steady state values for E_Cl_, E_K_ and V_m_ in our model showed that changing the average charge of impermeant anions, while substantially affecting E_Cl_, had very small effects on the driving force for Cl^−^ (Fig. 5B). By shifting z within reasonable ranges for mammalian neurons (Lodish et al., 2009; Raimondo et al., 2015), and assuming osmo- and electro- neutrality, only shifts of < 1 mV could be generated. In addition, although the absolute number of impermeant anions (moles) remained constant throughout the process of modifying z, cell volume shifted, and as a consequence modest alterations to the concentration of impermeant anions occurred as well.

**Figure 5:**
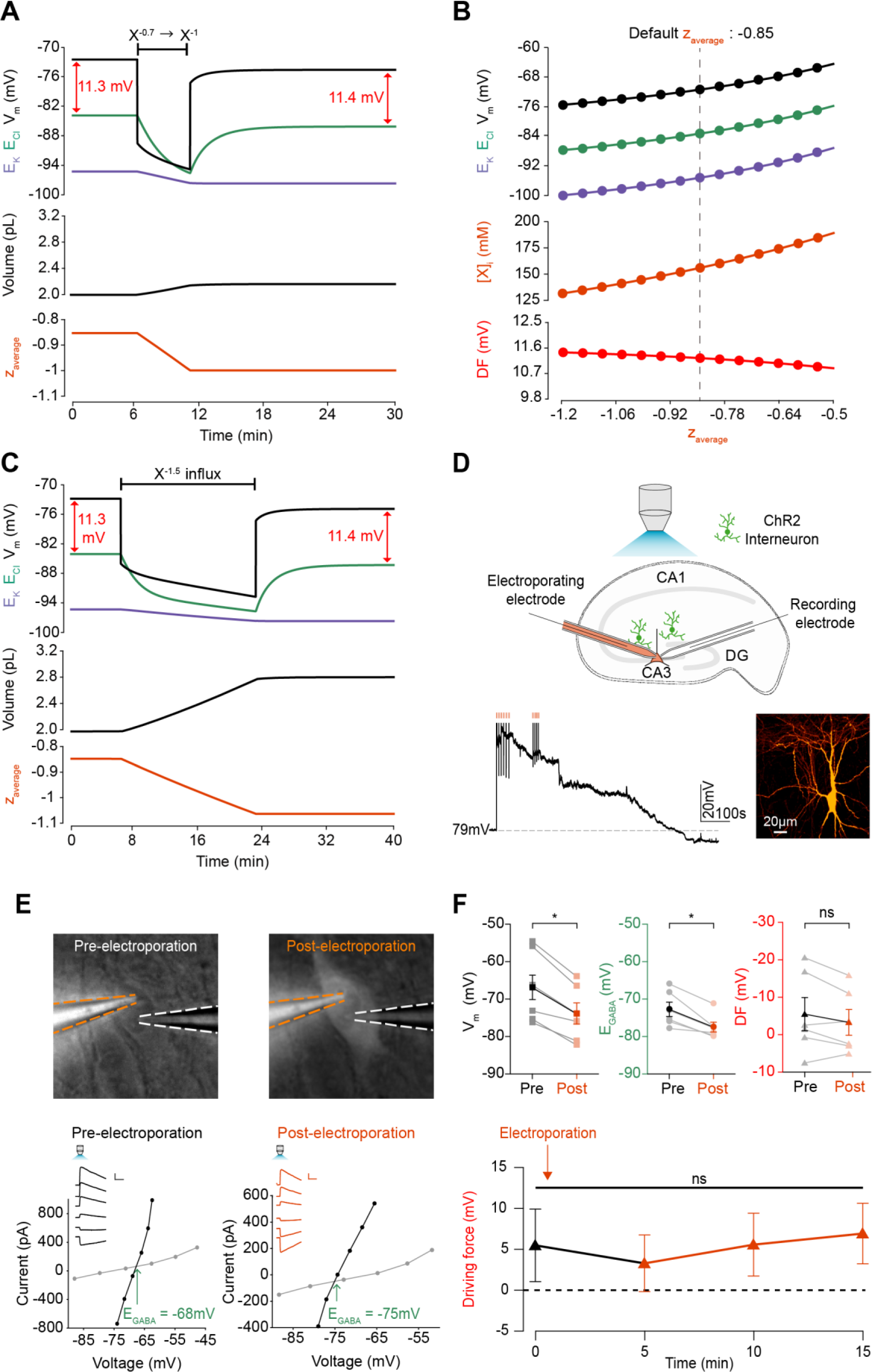
Adjusting the average charge of impermeant anions shifts the chloride reversal potential with negligible effects on the driving force for chloride. *(A) Decreasing the average charge of impermeant anions from −0.85 (the default value) to −1 (orange, bottom panel), without changing the absolute number of intracellular impermeant anions caused a persistent decrease in E_Cl_ (green, top panel), E_K_ (purple) and V_m_ with moderate increases in volume (middle panel) in our default single compartment model. Negligible changes in Cl^−^ driving force (ΔDF = 0.16 mV, red) were observed. (B) Analytic solution (solid lines) for different impermeant anion average charge (z) exactly matches the steady state values from numerical, time series runs (dots) based on adjusting average z as in ‘A’. Steady state E_Cl_, E_K_, V_m_ (top panel) and [X]_i_ (middle) increased with increasing average z, while changes in average z resulted in very small changes in Cl^−^ DF (bottom, red). The vertical dashed line indicates the values at the chosen default average z of −0.85. (C) Influx of a species of impermeant anions with average charge −1.5, that decreased average z (bottom panel) from −0.85 to −1 and increased the number of impermeant anions also caused persistent decreases of E_Cl_, E_K_ and V_m_ as in ‘A’, but with larger increases in cell volume (middle panel). Again, very small persistent changes in Cl^−^ DF were observed. The volume changes for different flux amounts and charges are illustrated in Figure 5 – figure supplement 1. (D) Top, schematic of the experimental setup where whole-cell recordings were made from CA3 pyramidal cells in mouse organotypic brain slices. Impermeant anions (orange) were delivered via electroporation of the negatively charged fluorescent dextran Alexa Flour 488 via a pipette positioned near the soma of the recorded cell. GABA_A_R currents were elicited via photo-activation (100 ms, 470 nm LED via objective) of ChR2-expressing GAD2+ interneurons (green cells) in the presence of 5 μM CGP-35348 to block GABA_B_Rs. Lower trace, current clamp recording showing V_m_ changes during electroporation of anionic dextran. Confocal image demonstrating cell-localised fluorescence of the anionic dextran electroporated in ‘E’. (E) Top, widefield images with electroporation pipette (orange dashed lines) and the recording pipette (white dashed lines). Note increased fluorescence in the soma after electroporation. Below, insets show GABA_A_R currents evoked by photo-activation of GAD2+ interneurons at different holding potentials. Calibration: 1 s, 100 pA. IV plots were used to calculate V_m_, E_GABA_ and DF before (left) and after (right) electroporation. (F) Top, population data showing significant decreases in mean V_m_ and E_GABA_ but not DF five minutes after electroporation. Below: changes in DF over time. Point at which electroporation occurred marked with orange arrow. ‘ns’, non-significant; *p < 0.05*.

Next, instead of adjusting the charge of existing anions in intracellular impermeant anions as described above, we directly added impermeant anions of differing valence to the cell (Fig. 5C and D). This had the effect of both increasing the absolute quantity of impermeant anions and adjusting the average charge of impermeant anions. The ‘addition’ of impermeant anions in this way models the *de novo* synthesis of impermeant anion species, or their active transport into the cell. This process also resulted in both transient and persistent changes to E_Cl_, E_K_ and V, which was dependent on the extent that average z was altered. Again, whilst the large additions of impermeant anions could substantially alter the Cl^−^ reversal potential, this had a negligible effect on the driving force for Cl^−^ due to matching shifts in V_m_. Driving shifts in E_Cl_ in this manner also resulted in changes to cell volume (**Figure 5 – figure supplement 1**).

To experimentally test our biophysical modelling predictions, we used photo-activation of ChR2 expressing GABAergic interneurons and whole-cell patch clamp recordings of mouse organotypic CA3 hippocampal pyramidal neurons to measure V_m_, E_GABA_, and GABA driving force (which approximates Cl^−^ driving force) before and after addition of impermeant anions (Fig. 5D). To add impermeant anions to the recorded cell, we used single cell electroporation of fluorescently tagged anionic dextrans (Alexa Flour 488, see Materials and Methods). Successful addition of impermeant anions could be confirmed visually by observing strong and stable fluorescence of the anionic dextran restricted to the recorded cell (Fig. 5D, E). To drive impermeant anions into the cell, negative voltage steps (20 ms, 0.5–1 V) were applied to the electroporation pipette necessarily resulting in direct membrane depolarization, which recovered over a period of 1-5 minutes (Fig. 5D). Once this acute perturbation had settled, as predicted by our model, changing the average charge of impermeant anions, following the addition of highly negatively charged dextrans to the cell, resulted in a stable, mean negative shift in V_m_ from a baseline of −66.95 ± 3.96 mV to −73.96 ± 3.06 mV (n = 6, *P* = 0.03, *Wilcoxon test*, Fig. 5E, F). Again in line with our predictions, addition of impermeant anions also resulted in a significant reduction of resting E_GABA_ (a proxy for E_Cl_) from baseline values of −72.45 ± 1.95 mV to −77.25 ± 1.35 mV (*P* = 0.03, *Wilcoxon test*). Importantly, however, similar shifts in V_m_ and E_GABA_ resulted in an undetectable shift in GABA and hence Cl^−^ driving force (5.49 ± 4.43 mV vs 3.33 ± 3.46 mV, n = 6, *P* = 0.22, *Wilcoxon test*, Fig. 5E, F).

Our single compartment model of Cl^−^ homeostasis, in conjunction with experimental validation, demonstrates that whilst the adjustment of average impermeant anion charge can significantly affect E_Cl_, this results in negligible changes to the driving force for Cl^−^. This contrasts with the results shown earlier, where adjusting the activity of cation-chloride cotransport modulates both E_Cl_ and the driving force for Cl^−^ substantially.

### Impermeant anions drive small shifts in chloride driving force by modifying the Na^±^/K^±^-ATPase pump rate under conditions of active chloride cotransport

We next set out to determine how, and under what conditions, the modification of impermeant anions could potentially generate the very small persistent shifts in Cl^−^ driving force we observed in our models. Due to their small size (< 1 mV) these were not detectable during the experimental validation. First, we repeated the simulation performed in Figure 5A by changing the average charge of impermeant anions in the cell, but under conditions where the Na^+^/K^+^-ATPase effective pump rate (J_p_) was either a cubic function of the transmembrane Na^+^ gradient (default condition) or was fixed at a constant value (Fig. 6A). In the case where the pump rate was fixed, adjusting the average charge of impermeant anions generated no persistent change in Cl^−^ driving force (Fig. 6A). Modifying impermeant anions caused a significant change in steady-state intracellular Na^+^ concentration when J_p_ was kept constant. However, small shifts in Cl^−^ driving force occurred only when the effective pump rate was variable, in which minor changes to [Na^+^]_i_ caused significant changes to J_p_, which in turn resulted in a small shift in Cl^−^ driving force. There is a direct relationship between the average charge of impermeant anions (z), [Na^+^]_i_, the effective Na^+^/K^+^-ATPase pump rate and Cl^−^ driving force. This relationship was abolished when the effective Na^+^/K^+^-ATPase pump rate was held constant by removing its dependence on Na^+^ (Fig. 6B). In addition, even large variations in effective pump rate near the default value caused negligible shifts in Cl^−^ driving force of < 1 mV. These results were similar when using the experimentally-matched ATPase model by Hamada et al. (2003), with slight differences in final values (**Figure 6 – figure supplement 1A**). These small, impermeant anion driven, Na^+^/K^+^-ATPase pump dependent shifts in Cl^−^ driving force are completely dependent on the presence of cation-chloride cotransport. In the absence of KCC2, there is no Cl^−^ driving force as E_Cl_ = V_m_ (Fig. 6F).

**Figure 6:**
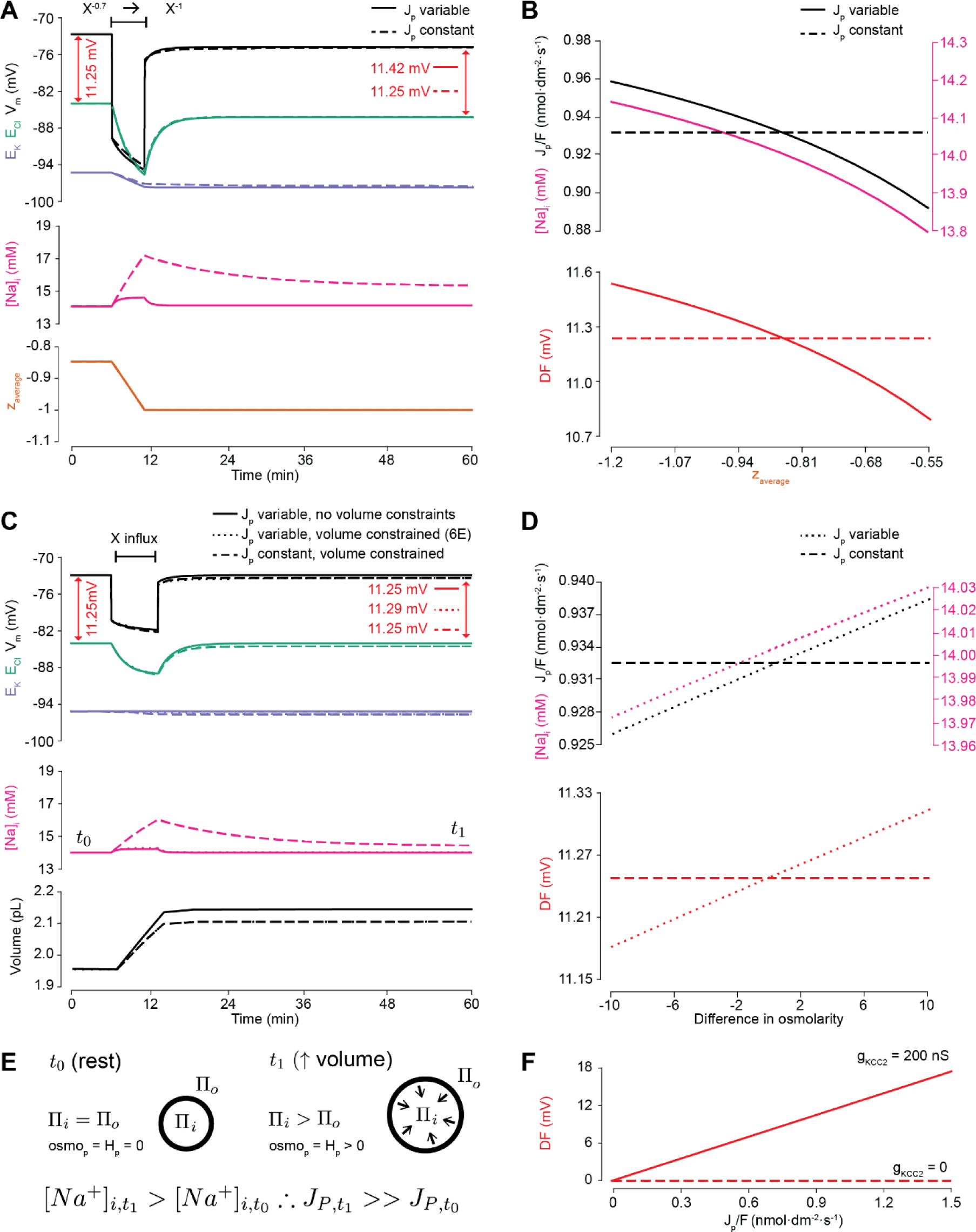
Impermeant anions drive small shifts in chloride driving force by modifying the Na^+^/K^+^-ATPase pump rate under conditions of active chloride cotransport. *(A) E_Cl_ (green), E_K_ (purple) and V_m_ (black) (top panel), [Na^+^]_i_ (pink, middle panel) and average impermeant anion charge z (orange, lower panel) over time in the default single compartment model. Changing the average z from −0.85 to −1 generated small, persistent Cl^−^ driving force (DF) shifts (arrows, red) only when the effective ATPase pump rate (J_p_) was variable (solid line) and not when J_p_ was kept constant (dashed line). (B) Solving analytically across different values of z with either a variable J_p_ (solid lines) or a constant J_p_ (dashed line), demonstrates the direct relationship between Na^+^ (pink), effective pump rate (J_p_) (top panel) and DF (lower panel, red). (C) E_Cl_ (green), E_K_ (purple) and V_m_ (black) (top panel), [Na^+^]_i_ (pink, middle panel) and cell volume (black, lower panel) over time in the default single compartment model. Impermeant anions of the same average charge of the cell were added. A volume constraint was incorporated by adding a hydrostatic force dependent on membrane tension (dashed lines), which resulted in an impermeable anion induced transmembrane osmotic differential. This caused a small change in DF when J_p_ was variable (dashed line), but not when J_p_ was held constant (dotted lines). (D) Solving analytically across osmolarity differences demonstrates the direct relationship between Na^+^ (pink), effective pump rate (J_p_) (top panel) and DF (lower panel). Note, the small changes in DF. (E) Schematic explaining the mechanism through which impermeant anion induced cell swelling in the presence of volume constraints (i.e. membrane tension) result in steady states with equal but non-zero osmotic (osmo_P_) and hydrostatic pressures (HP), causing transmembrane osmotic differences (t_1_). This causes small changes in Na^+^, and hence J_p_. (F) All Na^+^/K^+^-ATPase pump rate-related shifts in the DF require KCC2 activity; in the absence of activity (dashes), no shifts in driving force can occur. In Figure 6 – supplement figure 1, we show that the results in (A) and (C) are similar when an experimentally- matched model of the Na^+^/K^+^-ATPase is used*.

We then tested whether relaxing the condition of transmembrane osmoneutrality might also alter impermeant anion induced effects on Cl^−^ driving force. We modelled a situation where increases in cell surface area beyond a certain ‘resting’ surface area generated a hydrostatic pressure (membrane tension), which could balance an osmotic pressure difference of 10 mM between the intra and extracellular compartments (see Fig. 6C, schematic in Fig. 6E and Materials and Methods). In this case, adding impermeant anions of default charge z resulted in constrained increases in cell volume, which were accompanied by persistent transmembrane differences in osmolarity and intracellular Na^+^ concentration. This was sufficient to generate small differences in driving force for Cl^−^ of < 0.2 mV for reasonable increases in cell surface area (Nichol and Hutter, 1996; Dai et al., 1998). Again, this was entirely due to Na^+^ driven shifts in the Na^+^/K^+^-ATPase effective pump rate. By removing the dependence of Na^+^/K^+^-ATPase activity on Na^+^ concentration, addition of impermeant anions no longer generated persistent shifts in Cl^−^ driving force (Fig. 6C). Because the model is also directly dependent on [Na^+^]_i_, the results were similar when using the experimentally-matched ATPase model by Hamada et al. (2003) (**Figure 6 – figure supplement 1B**). We observed a direct relationship between transmembrane osmotic gradient, [Na^+^]_i_, the effective Na^+^/K^+^-ATPase pump rate and Cl^−^ driving force. This relationship was removed when the effective Na^+^/K^+^-ATPase pump rate was held constant, with no changes in Cl^−^ driving force seen despite the generation of the same shift in the transmembrane osmotic gradient (Fig. 6D).

In summary, changes in Cl^−^ driving force generated by changing the ionic contributions to cellular charge (by altering the average charge of impermeant anions) or osmoneutrality (by increasing the contribution of hydrostatic pressure) are due to the alteration of the dynamics of active ion transport mechanisms in the cell. However, these effects are negligible in magnitude and cannot contribute significantly to setting physiologically observed Cl^−^ driving forces. It is worth reiterating that any non-zero Cl^−^ driving force is entirely dependent on the presence of active Cl^−^ cotransport. In our model, in the absence of KCC2, neither the Na^+^/K^+^-ATPase nor impermeable anions can shift Cl^−^ out of equilibrium (Fig. 6F).

### Changes in cation-chloride cotransport activity generate local differences in chloride reversal and driving force, which depend on cytoplasmic diffusion rates

An important functional question is how Cl^−^ driving force might be modified at a local level within a neuron. We considered local persistent changes of Cl^−^ driving force for the case of active transmembrane Cl^−^ fluxes (Fig. 7) and impermeant anions (Fig. 8) by extending the single compartment model described above into a multi-compartment model or ‘virtual dendrite.’ This dendrite was 100 μm in length and consisted of ten compartments, each of 10 μm length and 1 μm diameter. The compartments contained the same mechanisms and default parameterization as the single compartment model described above. Compartmental volume was changed by altering the radius, while holding the length constant. In addition, all ions, except impermeant anions, could move between compartments by electrodiffusion (Fig. 7A and Materials and Methods).

**Figure 7:**
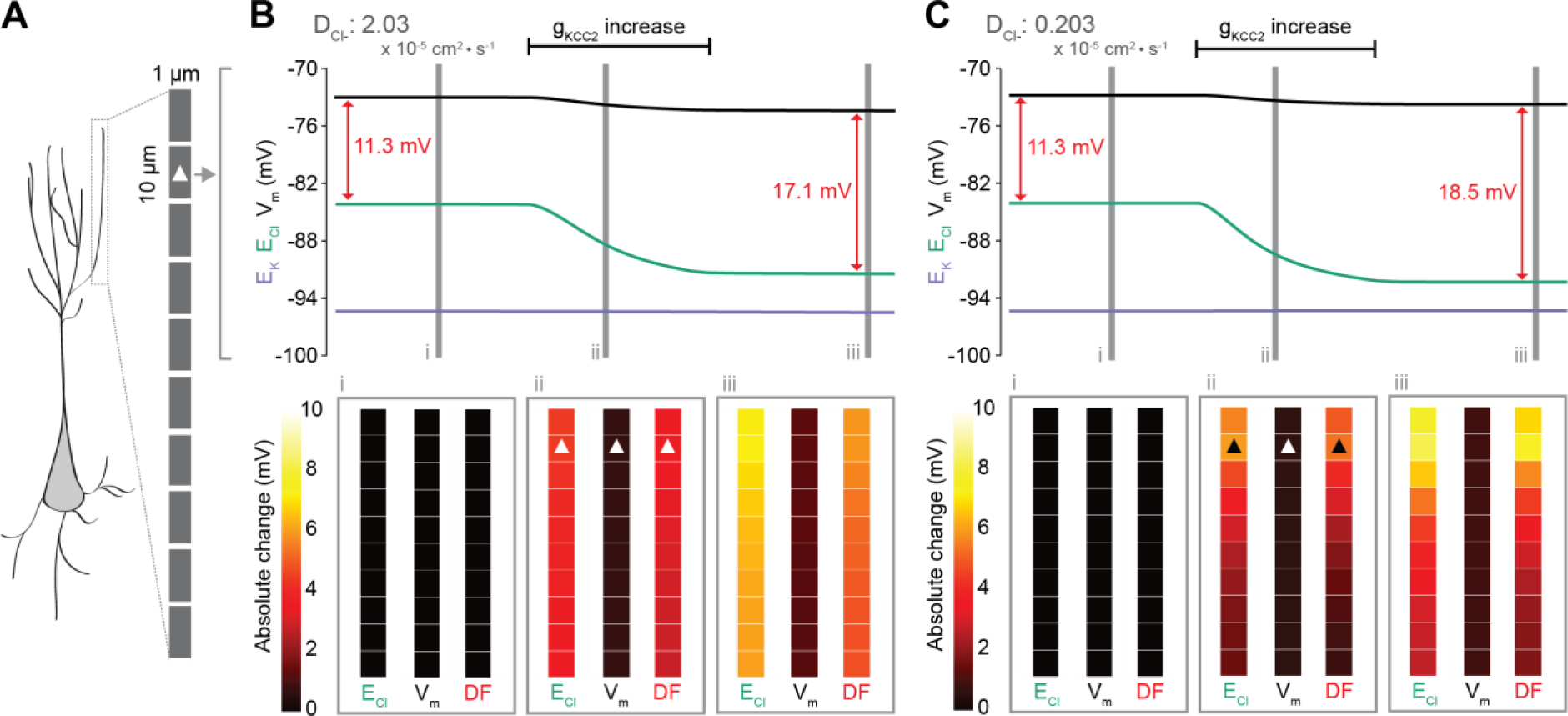
Local changes in KCC2 activity generate local differences in chloride reversal and driving force only under conditions of constrained chloride diffusion. *(A) Schematic depicting the multi-compartment model representing a virtual dendrite of length 100 μm. The virtual dendrite consists of 10 compartments of length 10 μm and initial radius 0.5 μm. Each compartment contains the same mechanisms and default parameterization as the single compartment model. All ions, except impermeant anions, could move between compartments by electrodiffusion. (B) Top panel, E_Cl_ (green), E_K_ (purple), V_m_ (black) and calculated DF (arrows, red) from the 2^nd^ from top compartment (indicated with a white triangle) where the conductance of KCC2 was increased. The insets depict the diameter, and absolute change from baseline of E_Cl_, V_m_ and DF for all compartments before (i), during (ii) and after (iii) the activity of KCC2 was selectively increased. This resulted in E_Cl_ decreasing in all compartments with minimal changes to E_K_ and V. Consequently, the Cl^−^ DF (red) increased. In this case the diffusion constant for Cl^−^ (D_Cl_ = 2.03 × 10^−7^ dm^2^.s^−1^) resulted in E_Cl_ and DF changes being widespread across the virtual dendrite. (C) Reducing the Cl^−^ diffusion constant to 0.2 × 10^−7^ dm^2^.s^−1^ resulted in a localized effect of compartment specific KCC2 activity increases on E_Cl_ and DF*.

**Figure 8:**
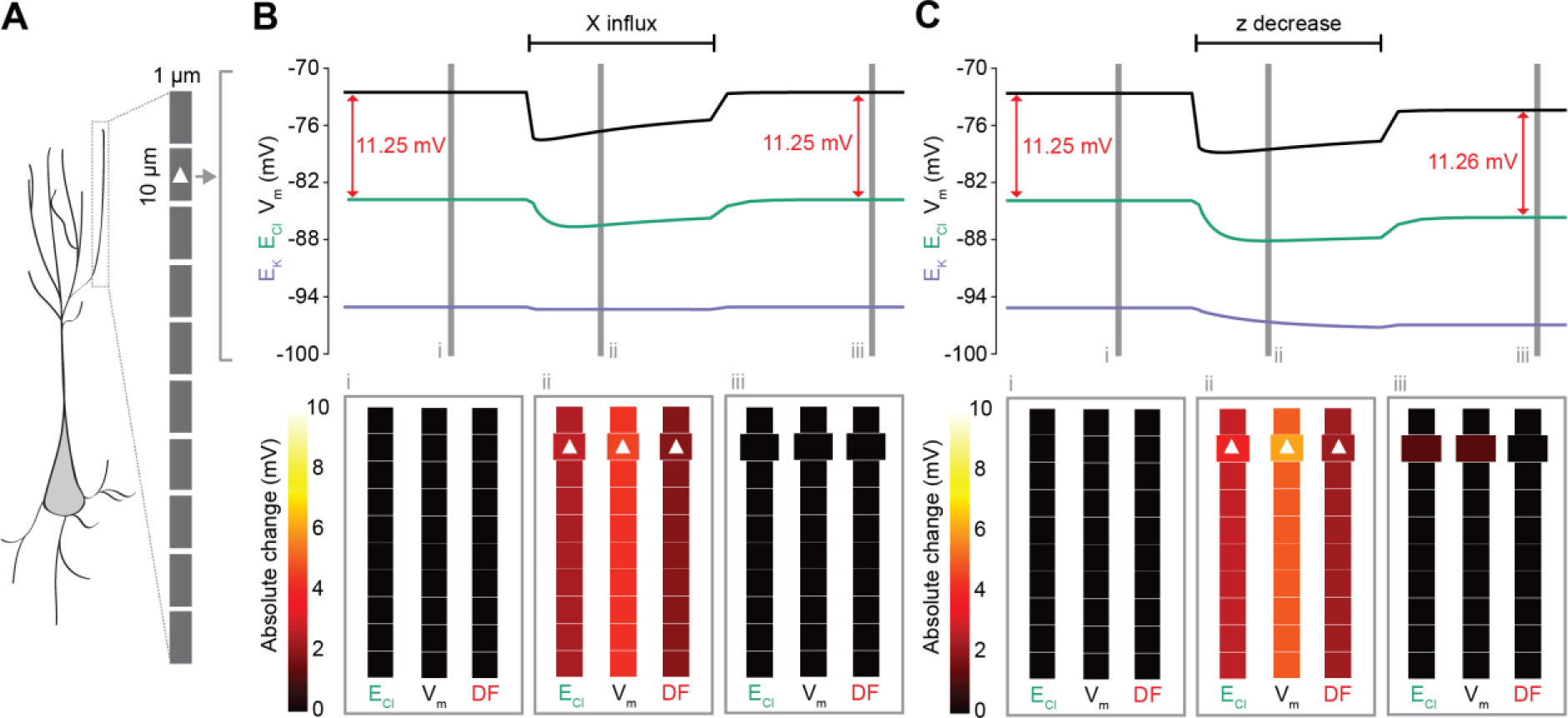
Local changes in impermeant anions do not establish the local driving force for chloride. *(A) Multi-compartment model of a 10 compartment 100 μm virtual dendrite as in Figure 7. (B) Top panel, E_Cl_ (green), E_K_ (purple), V_m_ (black) and calculated DF (arrows, red) within the compartment where additional impermeant anions were exclusively added (indicated with a white triangle). The insets depict the diameter, and absolute change from baseline for E_Cl_, V_m_ and DF for each compartment of the virtual dendrite before (i), during (ii) and after (iii) impermeant anions were added. The selective addition of impermeant anions of average z to the 2^nd^ from top compartment resulted in transient but non-permanent shifts in E_Cl_, V_m_ and the Cl^−^ DF in all compartments. The volume of the compartment where impermeant anion addition occurred increased permanently. (C) Traces and insets as in ‘B’ showing the addition of impermeant anions of different charge in order to decrease average z in the 2^nd^ from top compartment specifically. Note that during addition of impermeant anions, E_Cl_, E_K_, V_m_ changed. We also observed persistent decreases in E_Cl_ and V_m_ in the compartment manipulated, with a negligible change in DF (0.01 mV). Again, impermeant anion addition resulted in an increase in the volume of the specific dendritic compartment manipulated*.

To explore the local effects of CCC activity, we increased g_KCC2_ from our default value of 20 µS/cm^2^ to 600 µS/cm^2^ in the second last compartment of the virtual dendrite exclusively. This resulted in a persistent decrease in E_Cl_, concurrent with a modest decrease in V_m_, resulting in a permanent increase in Cl^−^ driving force and minimal change in compartment volume (Fig. 7B). The spatial precision of this alteration depended strongly on the diffusion constant for Cl^−^. With a Cl^−^ diffusion constant of 2.03 × 10^−7^ dm^2^.s^−1^, these alterations spread widely through the virtual dendrite. For example, the change in Cl^−^ driving force was 4.8 mV in the furthermost compartment (90 μm apart) as compared to 5.9 mV in the compartment manipulated. When we decreased the Cl^−^ diffusion constant by one order of magnitude, the change in Cl^−^ driving force was 7.3 mV in the compartment in which KCC2 was adjusted, but only 1.8 mV in the furthermost compartment from the site of manipulation (Fig. 7C). These findings suggest that local differences in cation-chloride cotransport activity can drive spatially restricted differences in Cl^−^ driving force under conditions of constrained Cl^−^ diffusion, however under conditions of typical ionic diffusion the effect of Cl^−^ transport by KCC2 is relatively widespread.

### Local impermeant anions do not appreciably affect the local driving force for chloride

Following the last result, we considered whether changing impermeant anions in part of a dendrite could create a local area with a different Cl^−^ driving force compared to the rest of the cell. We first added impermeant anions of average charge (z) exclusively to the second-last compartment of the virtual dendrite while measuring the Cl^−^ reversal, V_m_ and Cl^−^ driving force in all compartments. During addition of the impermeant anions, E_Cl_ and V_m_ decreased with an accompanying decrease in Cl^−^ driving force. However, following cessation of impermeant anion influx, all parameters returned to baseline levels, except for the volume of that specific compartment, which showed a modest increase (Fig. 8B). This suggests that local addition of impermeant anions of average charge has no local effect on Cl^−^ homeostasis but can affect the volume of the compartment concerned.

Next, we again added impermeant anions to the second-last compartment of the virtual dendrite, but this time we added impermeant anions with a more negative charge than average (z = −1). This resulted in the average charge of impermeant anions in that specific compartment becoming more negative (Fig. 8C). During the addition of the impermeant anions, E_Cl_ and V_m_ decreased across the dendrite, but with small accompanying shifts in Cl^−^ driving force. Following cessation of local impermeant anion influx, a persistent shift in E_Cl_ and V_m_ was observed specifically in the compartment manipulated. However, this generated a negligible, persistent change in Cl^−^ driving force (< 0.01 mV change in driving force for a compartment specific change in z from −0.85 to −0.93), and only within that specific compartment of the virtual dendrite. Again, impermeant anion addition resulted in a permanent increase in the volume of the compartment concerned. This finding suggests that local impermeant anions can adjust the Cl^−^ reversal potential locally, but are not well-placed to cause significant, permanent shifts in the driving force for Cl^−^. Indeed, electrodiffusion may further limit the degree to which changes in impermeant ion charge can modify driving forces through alterations in active ionic transport: the resulting permanent Cl^−^ driving force changes in the multi-compartment model are many times smaller than the shifts in the single compartment version (as compared to Fig. 6B).

## Discussion

The driving force for Cl^−^ is a fundamental parameter affecting the excitability of neuronal networks (Raimondo et al., 2017). Recently, impermeant anions, rather than CCCs, have been suggested as the primary determinants of the neuronal driving force for Cl^−^ (Glykys et al., 2014). Here we have explored the determinants of the Cl^−^ driving force in neurons by deriving theoretical models based on biophysical first principles. We show that the Na^+^/K^+^-ATPase, baseline K^+^, Na^+^ and Cl^−^ conductances, average charge on impermeant anions, water permeability and CCCs, likely all play roles in setting neuronal [Cl^−^]_i_. However, our findings suggest that while impermeant anions can contribute to setting the [Cl^−^]_i_ in neurons, they can only affect Cl^−^ driving force by modifying the activity of active transport mechanisms (i.e. the Na^+^/K^+^-ATPase). Our modelling and experimental data demonstrate that under physiologically relevant conditions, impermeant anions do not alter the Cl^−^ driving force significantly. In contrast, CCCs are well placed to modulate Cl^−^ driving force and hence inhibitory signaling.

Previous theoretical models, which account for the dynamics of Cl^−^ ions, have been useful in determining how changes to the driving force for Cl^−^ are critical for controlling the effect of synaptic inhibition in the brain (Qian and Sejnowski, 1990; Staley and Proctor, 1999; Doyon et al., 2011; Jedlicka et al., 2011; Lewin et al., 2012; Mohapatra et al., 2016). Whilst these models have included the Na^+^/K^+^-ATPase, the interacting dynamics of several ion species, CCCs (Doyon et al., 2011; Krishnan and Bazhenov, 2011), electrodiffusion (Qian and Sejnowski, 1989) and impermeant anions and volume regulation (Dijkstra et al., 2016), none have combined all of these mechanisms to explore how their combination determines the local driving force for Cl^−^. Our theoretical approach is based on the pump-leak formulation (Tosteson and Hoffman, 1960). It suggests that mammalian cells maintain their volume under osmotic stress generated by impermeant anions and the Donnan effect by employing active transport of Na^+^ and K^+^ using the Na^+^/K^+^-ATPase (Armstrong, 2003; Kay, 2017). A Donnan equilibrium, a true thermodynamic equilibrium requiring no energy to maintain it, is not possible in cells with pliant membranes like neurons (Sperelakis, 2012).

Our model conforms to the pump-leak formulation: abolishing the activity of the Na^+^/K^+^-ATPase leads to cell swelling, progressive membrane depolarization and rundown of ionic gradients, including that of Cl^−^. Therefore, the Na^+^/K^+^ ATPase is a fundamental cellular parameter that stabilizes cell volume and determines all ionic gradients including that of Cl^−^ and hence must be considered in any attempt to model ion homeostasis. Interestingly however, we demonstrate that above a certain level of Na^+^/K^+^-ATPase activity, even many fold changes in pump rate have minimal effects on volume, E_Cl_ and V_m_. This might explain recent experimental findings in which periods of Na^+^/K^+^ ATPase inhibition using ouabain caused modest changes to cell volume (Glykys et al., 2014). It therefore seems unlikely that neurons adjust the Na^+^/K^+^-ATPase as a means for modulating Cl^−^ driving force.

Baseline ion conductances are another important factor that affect Cl^−^ driving force. Our model is consistent with recent experimental results that demonstrate that increased neuronal Na^+^ conductance (for example by activating NMDA receptors, or preventing closure of voltage-gated Na^+^ channels), leads to progressive neuronal swelling, membrane depolarization and Cl^−^ accumulation (Rungta et al., 2015) – the primary pathological process in cytotoxic oedema (Liang et al., 2007). We also show that tonic neuronal Cl^−^ conductance only affects baseline [Cl^−^]_i_ and driving force in the presence of CCCs. Without active Cl^−^ flux, which CCCs provide, there is no driving force for passive Cl^−^ flux and hence no mechanism for [Cl^−^]_i_ changes resulting from selective modification of a Cl^−^ conductance. This is consistent with both classic (Misgeld et al., 1986; Thompson and Gähwiler, 1989) and recent experimental findings (Berglund et al., 2016).

In our model, we find that elevating the activity of KCC2, the most active CCC in mature neurons (Ben-Ari, 2002), increases the driving force for Cl^−^ by shifting the reversal potential for Cl^−^ closer to that of K^+^. Interestingly, large shifts (∼7 mV) in driving force were associated with very minor (1%) changes in volume or membrane potential. As such, modulating KCC2 represents a specific means for manipulating the neuronal Cl^−^ driving force. This is consistent with traditional dogma, recent and previous experimental results (Kaila et al., 2014; Klein et al., 2017) as well as our own experimental validation using furosemide to block the activity of KCC2, which drove significant changes in driving force with little effect on V_m_. In further support of this, our meta-analysis of numerous experimental studies showed a strong correlation between change in KCC2 expression and Cl^−^ driving force, but not between KCC2 expression and V_m_. There is an ongoing debate as to whether some cotransporters, including CCCs, might also couple water transport to ion transport (Zeuthen, 1994; MacAulay et al., 2002; Gagnon et al., 2004; Charron et al., 2006). Although we did not model the active movement of water by KCC2, this scenario would not alter the central importance of CCCs for setting the Cl^−^ driving force.

Using our multi-compartment model, which incorporated electrodiffusion, we found that local modification of KCC2 activity has a specific local effect on Cl^−^ driving force that is dependent on the characteristics of intracellular Cl^−^ diffusion. Cytoplasmic Cl^−^ diffusion rates had to be reduced substantially before we observed local changes in Cl^−^ driving force driven by KCC2 (Qian and Sejnowski, 1989; Kuner and Augustine, 2000). Whilst differences in KCC2 activity might generate a gradient in Cl^−^ driving force between large subcellular structures (i.e. dendrites versus soma), our modelling results call into question the idea of synapse-specific regulation of Cl^−^ driving force within the same cellular domain (Földy et al., 2010).

Glykys et al. (2014) used Cl^−^ imaging and various experimental manipulations to claim that [X]_i_ and [X]_o_ sets [Cl^−^]_i_ and the Cl^−^ driving force. From our theoretical analysis, we find that modifying the amount of impermeant anions inside or outside neurons has no persistent effect on [Cl^−^]_i_ or Cl^−^ driving force, unless we include a mechanism that allows a transmembrane osmotic pressure differential to develop that indirectly affects active transport mechanisms. Even in this case, under transmembrane pressure differentials that do not lyse the membrane (Nichol and Hutter, 1996), Cl^−^ driving force changes are negligible (< 1 mV). Recently it has been suggested that the viscoelastic properties of the cellular cytoskeleton could allow it to take up osmotic shifts created by impermeant anion movement like a sponge (Sachs and Sivaselvan, 2015). This would mean that one would not see as large a volume shifts as predicted by our models. In our model we have assumed that water can pass through the neuronal membrane to equalize osmotic differences. Although it is thought that some neurons do not express aquaporin channels (Andrew et al., 2007), water can permeate the phospholipid bilayer (Fettiplace and Haydon, 1980). Therefore, whilst differences in neuronal water permeability might affect the time taken to reach steady-state, the steady state values themselves are unlikely to be affected. We conclude that [Cl^−^]_i_ and the Cl^−^ driving force are not determined by the *concentration* of impermeant anions.

However, our theoretical findings offer a potential explanation for recent experimental observations. We show that modifying the average *charge* upon impermeant anions (i.e. z in [X^z^]_i_), rather than their concentration, can affect [Cl^−^]_i_ and E_Cl−_. Relating this to prior experimental observations, Glykys et al. (2014) used SYTO64 staining of nucleic acids and perfusion of weak organic acids in conjunction with Cl^−^ imaging to suggest that [Cl^−^]_i_ depends upon internal impermeant anions ([X]_i_). If such a manipulation modifies the average charge on internal impermeant anions, and not concentration *per se*, this could account for the observed changes in [Cl^−^]_i_. Glykys et al. (2014) did not measure V_m_ or the Cl^−^ driving force in these experiments. The clear prediction from our model is that any manipulation, which changes the average charge of impermeant anions would not appreciably affect the Cl^−^ driving force because any impermeant anion driven change on E_Cl−_ is matched by an equivalent effect on V_m_ due to accompanying shifts in cation concentrations. We have provided experimental support for this prediction by showing that whilst E_GABA_ (and E_Cl_) can be shifted by addition of impermeant anions using electroporation of membrane impermeant anionic dextrans, V_m_ is shifted in a similar direction resulting in an undetectable change in Cl^−^ driving force.

Given prior theoretical predictions (Kaila et al., 2014; Voipio et al., 2014; Savtchenko et al., 2017), it is interesting that our model reveals that changing impermeant anions could affect the Cl^−^ driving force at all. We found that the small (< 1 mV) impermeant anion-driven changes in Cl^−^ driving force observed in our model were caused by indirect effects on Na^+^ concentration and hence Na^+^/K^+^-ATPase activity. The impermeant anion-driven changes in Cl^−^ driving force are even smaller in the multi-compartment model (< 0.1 mV), in which electrodiffusion allows local changes in Na^+^ to dissipate. When Na^+^/K^+^-ATPase activity was decoupled from the transmembrane Na^+^ gradient, we found that impermeant anions were unable to cause persistent shifts in Cl^−^ driving force as predicted theoretically (Kaila et al., 2014; Voipio et al., 2014; Savtchenko et al., 2017). It is important to note that these small, impermeant anion-Na^+^/K^+^-ATPase-driven shifts in Cl^−^ driving force are dependent on the presence of cation-chloride cotransport in the form of KCC2 and would entail changes in energy use by the Na^+^/K^+^-ATPase. In other words, active transport mechanisms are again required to drive changes in Cl^−^ homeostasis.

In summary, our theoretical models, which are derived from well-established physical principles, are consistent with our own experimental data and that of others (Glykys et al., 2014; Kaila et al., 2014; Klein et al., 2017), and suggest that impermeant anions alone cannot shift Cl^−^ out of equilibrium across the neuronal membrane. Were neurons to alter impermeant anion concentrations or charge, the resting membrane potential would be modified with little effect on the Cl^−^ driving force. Our work confirms the central importance of CCC activity in determining the effects of inhibitory synaptic transmission in the nervous system.

## Materials and methods

### Single compartment model

The single compartment model consisted of a cylindrical semi-permeable membrane separating the extracellular solution from the intracellular milieu with variable volume (Fig. 1A). The extracellular ionic concentrations were assumed constant (Table 1). Permeable ions in the model were K^+^, Na^+^ and Cl^−^ with their usual charges, while impermeant anions X were assumed to be a heterogeneous group of impermeant chemical species with average intracellular charge z and extracellular charge −1. z was chosen on the basis of known resting intracellular ion concentrations (Lodish et al., 2009; Raimondo et al., 2015) and osmolarity (Π). Bicarbonate ions were not included in our model as a permeant anion as they were assumed to be important for acute depolarizing effects (via GABA_A_Rs) rather than the chronic shifts in Cl^−^ driving force, which are the focus of this work (Staley and Proctor, 1999). The model included ionic leak currents for the permeable ions, Na^+^/K^+^-ATPase transporters and a CCC, in this case the K^+^-Cl^−^ cotransporter (KCC2). KCC2 and not NKCC1 is thought to be the most active CCC in mature neurons (Ben-Ari, 2002), therefore, to maintain conceptual simplicity only KCC2 was modelled. Cell volume (w) change was based on osmotic water flux and incorporated a membrane surface area scaling mechanism. An analytical solution to the model at steady state was derived using standard techniques and can be found in Supplementary file 1. The numerical model was initialised assuming conditions close to electroneutrality and an osmotic equilibrium between the intracellular and extracellular compartments. A forward Euler approach was used to update variables at each time step (dt) of 1 ms. Using a smaller dt did not influence the results in Figure 1–5. Code was written in Python 2 and is available on GitHub (https://github.com/kiradust/model-of-neuronal-chloride-homeostasis). The GitHub repository includes a file of Supplementary figures, in which we display transmembrane fluxes of all ions and water for relevant simulations. An example figure displaying ionic flux for all ions is available for Figure 4C in Figure 4 – figure supplement 1.

**Table 1:**
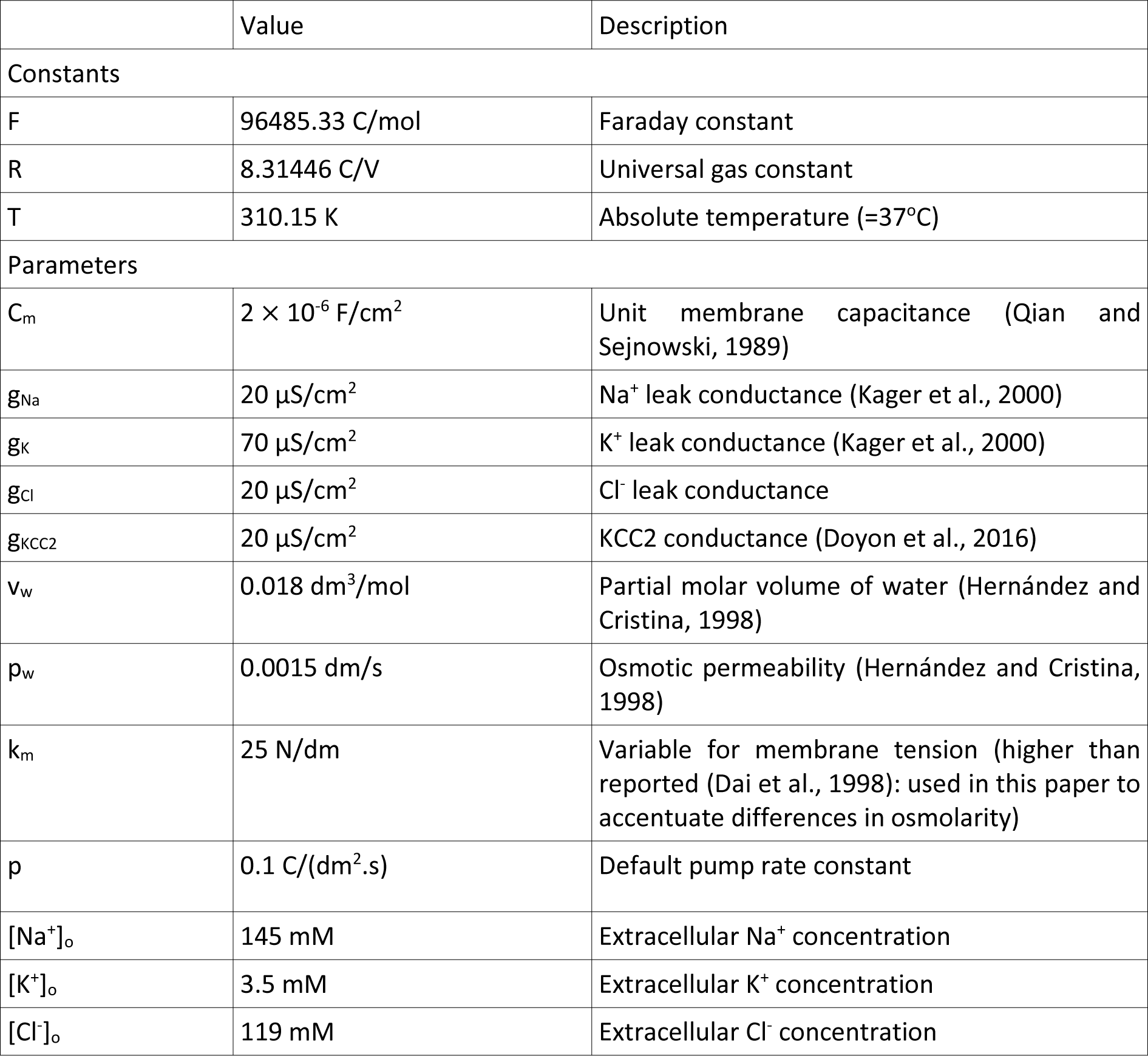
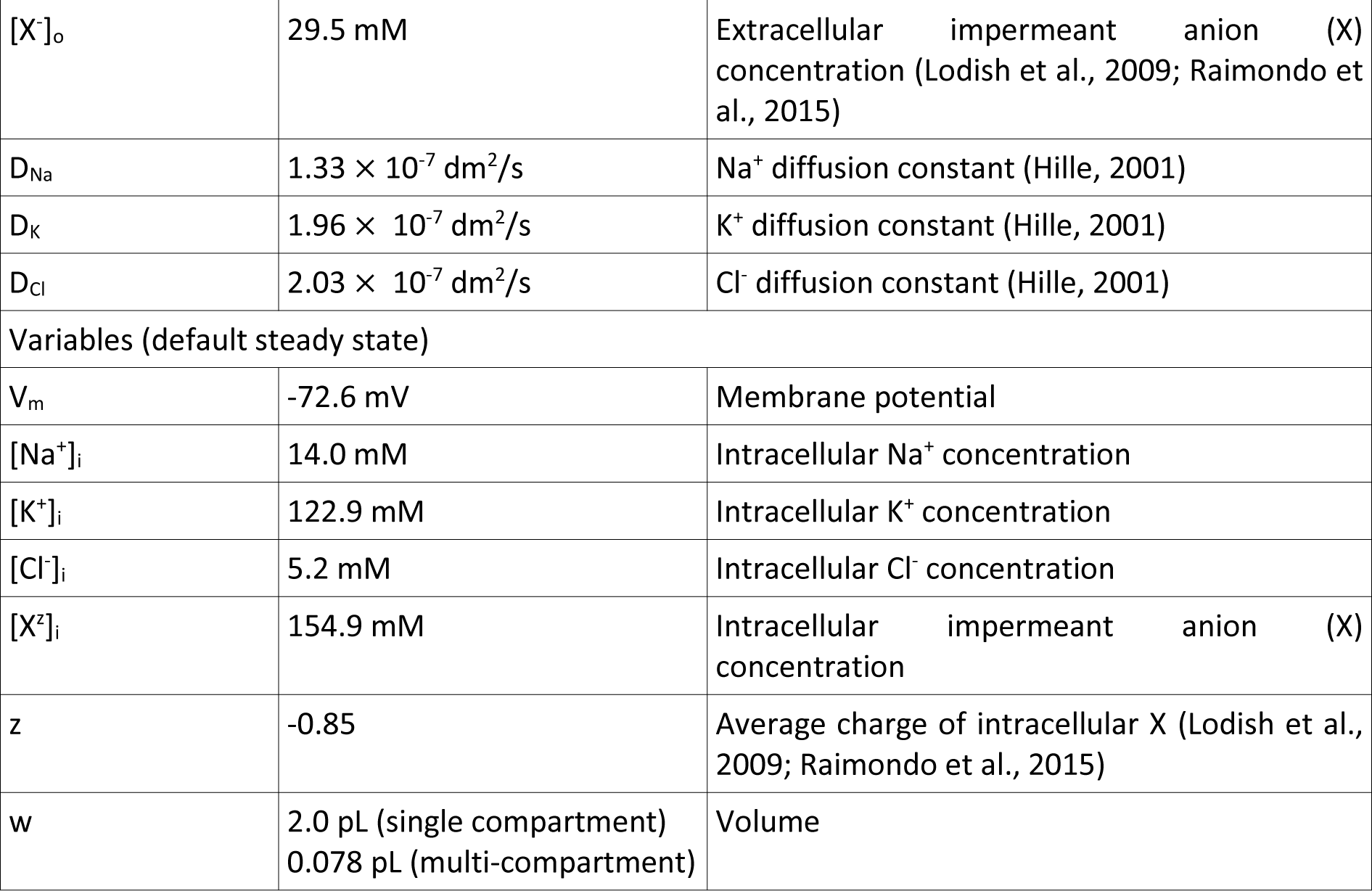
Constants, default parameters and usual steady state values for variables used in the biophysical models.

### Membrane potential

The membrane potential V_m_ was based on the “Charge Difference” approach of Fraser and Huang (2004) as follows:

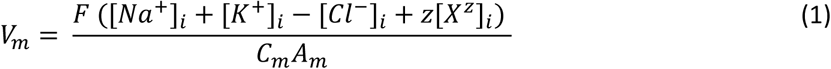

where F is Faraday’s constant, C_m_ is the unit membrane capacitance and A_m_ is calculated as the ratio of the surface area (of the cylinder) to cell volume. The term in brackets is the sum of all ionic charges within the cell. This approach has the advantage that the initial voltage can be calculated without needing to assume a steady state as is required for by the Goldman-Hodgkin-Katz (GHK) equation.

### Permeable ion concentrations

Intracellular concentrations of the permeable ions Na^+^, K^+^ and Cl^−^ were updated individually by summing trans-membrane fluxes. Leak currents were calculated using the standard equivalent circuit formulation *I* = *g*(*V_m_* − *E*). In addition, Na^+^ and K^+^ were transported actively by the Na^+^/K^+^-ATPase, with pump rate J_p_, which was approximated by a cubic function dependent on the transmembrane sodium gradient, following Keener and Sneyd (1998):

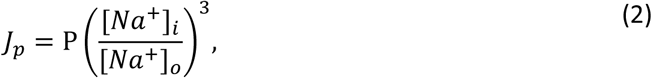

where P is the pump rate constant. Because it is a function of the sodium gradient, J_p_ decreases as [Na^+^]_i_ depletes. This formulation has been shown to be similar to more accurate kinetic models reliant on both the Na^+^ gradient and ATP concentration (Keener and Sneyd, 1998). To switch the ATPase pump on or off (Fig. 1C), P was decreased/increased exponentially over 10-20 minutes, consistent with previous reports of the dynamics of inhibition of the ATPase by ouabain and in turn the inhibition of ouabain’s effects by potassium canrenoate (Baker and Willis, 1972; Yeh and Lazzara, 1973). The ATPase pumps 2 K^+^ ions into the cell for every 3 Na^+^ ions out and these constants must be multiplied by J_p_ for each ion respectively. K^+^ and Cl^−^ were also modified by flux through the type 2 K-Cl cotransporter (KCC2), J_KCC2_ (Doyon et al., 2016):

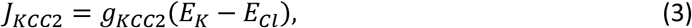

where g_KCC2_ is a fixed conductance and E_K_ and E_Cl_ are the Nernst potentials for K^+^ and Cl^−^ respectively. J_KCC2_ is 0 when E_K_=E_Cl_. The rate of change of the intracellular concentration of the three permeant ions was given by the following equations, with the Nernst potentials for each ion given by *E_ion_* = 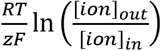, w indicating the cell volume, and 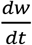 as described in equation (7) (Fig. 1–5, 6A, B, 7 and 8) or (9) (Fig. 6C-E):

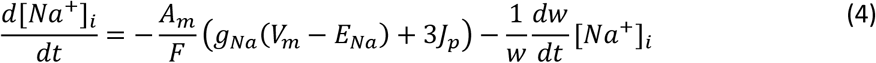

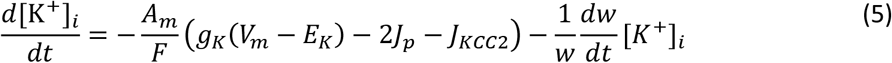

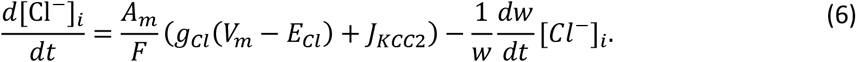

### Volume

Because the osmotic flux of water is expected to be faster than ion fluxes, the volume of the cell was adjusted to preserve transmembrane osmotic balance at each time step. Change in compartment volume, w, was calculated by changing the previous volume proportional to the difference between intracellular and extracellular osmolarity, Π_*i*_ and Π_*o*_ respectively (Hernández and Cristina, 1998), where v_w_ is the partial molar volume of water, p_w_ the osmotic permeability of a biological membrane and SA the surface area:

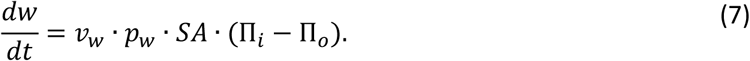

For the calculations in Figure 6C-E, where we allowed transmembrane differences in osmolarity to develop, we assumed that at rest the cylindrical cell had a radius of r_a_ and zero pressure across the membrane, and that the tension (T) in the membrane followed Hooke’s law such that the tension was proportional to the difference between the dynamic circumference of the cell and that of the resting state. From Laplace’s law the hydrostatic pressure in the cell was given by:

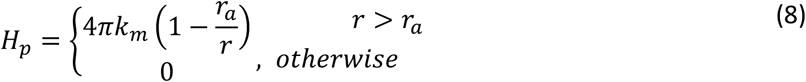

where k_m_ is the spring constant of the membrane (Sachs and Sivaselvan, 2015). Equation (7) was thus reformulated:

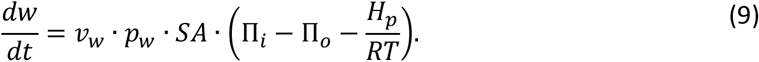

In order to simulate extreme conditions of constrained volume, a larger k_m_ was employed than is realistic (Dai et al., 1998). Intracellular ion concentrations were updated again after volume change at each time step. Volume changes were manifested in the cylindrical compartment as change in the radius. In Figure 1–6, the cell was initialised with diameter 10 μm and length 25 μm.

### Anion flux

Impermeant anions were manipulated in the compartment in Figure 4–6 and Figure 8 through several mechanisms. Anions could be added to the compartment at a constant rate dependent on A_m_ and have either the same average intracellular X charge z = −0.85 (Fig. 4C, 6C and 8B), or a different charge (Fig. 5C, 6A and 8C). In these cases, the number of moles of X in the compartment was increased. Alternatively, the charge of a species of intracellular X was slowly changed imitating a charge-carrying trans-membrane reaction (Fig. 5A and 6A). In this case, the number of moles of intracellular X did not change and it was assumed charge imbalance was mopped up by the extracellular milieu. Finally, extracellular X^−^ was changed in Figure 4D by removing as much Cl^−^ as X^−^ was added, thus maintaining osmolarity and electroneutrality in the extracellular space.

### Multi-compartment model

The single compartment dendrite model was incorporated in a multi-compartment model by allowing electrodiffusion to occur between individual compartments operating as described above. Compartments were initialised with a radius of 0.5 μm and length of 10 μm. Compartments were linked linearly without branching; 10 connected compartments in total were used. The time step dt was decreased to 10^−3^ ms for simulations in multiple compartments. Code was written in Python 3 and is available on GitHub (https://github.com/kiradust/model-of-neuronal-chloride-homeostasis).

### Electrodiffusion

The Nernst-Planck equation (NPE) was used to model one-dimensional electrodiffusion, based on Qian and Sejnowski (1989). The NPE incorporates fluxes because of diffusion and drift (i.e. the movement of ions driven by an electric field). It has been shown to be more accurate than using J_diffusion_ alone in small structures like dendrites (Qian and Sejnowski, 1989). The NPE for J the flux density of ion C is calculated as:

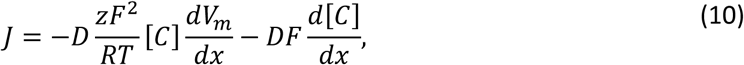

where D is the diffusion constant of ion C (Table 1), z is its charge, [C] is its concentration and x is the distance along the longitudinal axis over which electrodiffusion occurs. The NPE was implemented between compartments i and i+1, assuming the i→i+1 direction was positive, using a forward Euler approach. The midpoints of the compartments were used to calculate dx, i.e. 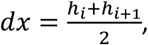 and the concentrations of C in each compartment were averaged to obtain J_drift_, ensuring that J_i→i+1_ = J_i+1→i_, where the fluxes had units of mol/(s.dm^2^):

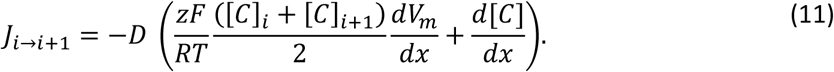

The flux was multiplied by the surface area between compartments and then divided by compartment volume to determine the flux in terms of molar concentration (M/s), i.e. 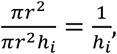 and finally implemented numerically with a forward Euler approach. The implementation mirrored that in Qian and Sejnowski (1989) for non-branching dendrites, but was adjusted for compartments whose volumes can change:

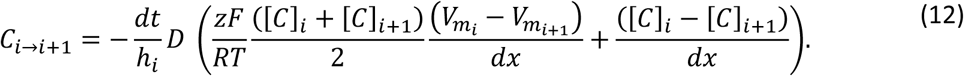

### Systematic review

A literature search was performed to identify experimental studies that aimed to correlate a change in KCC2 expression with changes in [Cl^−^]_i_. The MEDLINE database was used and accessed via the PubMed online platform. Search terms included ‘chloride’, ‘Cl’, ‘intracellular’, ‘KCC2’, ‘cotransporter’, ‘neuronal’, ‘GABA’ using appropriate Boolean operators. All 26 studies that demonstrated changes in KCC2 expression and E_GABA_ were considered for the meta-analysis. As there is a well described differential expression of KCC2 and NKCC1 at different stages of development, with KCC2 expression increasing and NKCC1 expression decreasing across development, only studies that used tissue older than postnatal day 7 were included (8 included data from younger animals). Other exclusion criteria included: reporting a significant change in NKCC1 (5 studies); use of non-rodent tissue (2 studies); no quantification of the change in KCC2 (2 studies). Nine experiments from eight studies met all criteria and were included (Coull et al., 2003; Lagostena et al., 2010; Lee et al., 2011; Ferrini et al., 2013; Campbell et al., 2015; MacKenzie and Maguire, 2015; Mahadevan et al., 2015; Tang et al., 2015). However, one study did not report the change in V_m_ and hence was excluded from the figure (Mahadevan et al., 2015). Data used in regression can be seen in Supplementary file 2 (Table S2-1) and includes the analysis for regression against change in [Cl^−^]_i_. To accommodate varied experimental preparations and techniques influencing data quality and biases, a 34-point scoring system was designed to weight the studies (Table S2-2). A weighted least squares regression model was then used to correlate the percentage (%) change in KCC2 expression versus change in Cl^−^ driving force.

### Slice preparation and electrophysiology

For all experiments, organotypic slices were prepared from rodent brain tissue. Wistar rats were used for the experiments testing the effects of CCC blockade. However, to allow for optogenetic manipulation, a crossed mouse strain on a C57BL/6 background was used. This mouse strain was a cross between mice expressing cre-recombinase in glutamic acid decarboxylase 2 (GAD2) positive interneurons (GAD2-IRES-cre, Jax Lab strain 010802) and mice with a loxP-flanked STOP cassette preventing transcription of the red-fluorescent protein tdTomato (a cre-reporter strain, Ai4, Jax Lab strain 007914.). This created the GAD2-cre-tdTomato strain which resulted in cre-recombinase and tdTomato expression in all GABAergic interneurons (Taniguchi et al., 2011). The use of all animals was approved by either the University of Oxford (rat) or the University of Cape Town (mouse) animal ethics committees.

Organotypic brain slices were prepared using 7 day old Wistar rats (CCC experiment) or crossed GAD2- IRES-cre (Jax Lab strain 010802) and Ai14 tdtomato reporter (Jax Lab strain 007914) mice (impermeant anion experiment) and followed the protocol originally described by (Stoppini et al., 1991). Briefly, brains were extracted and swiftly placed in cold (4°C) dissection media consisting of Gey’s Balanced Salt Solution (GBSS #G9779, Sigma-Aldrich, USA) supplemented with D-glucose (#G5767, Sigma-Aldrich, USA). The hemispheres were separated and individual hippocampi were removed and immediately cut into 350 μm slices using a Mcllwain tissue chopper (Mickle, UK). Cold dissection media was used to rinse the slices before placing them onto Millicell-CM membranes (#Z354988, Sigma-Aldrich, USA). Slices were maintained in culture medium consisting of 25% (vol/vol) Earle’s balanced salt solution (#E2888, Sigma-Aldrich, USA); 49% (vol/vol) minimum essential medium (#M2279, Sigma-Aldrich, USA); 25% (vol/vol) heat-inactivated horse serum (#H1138, Sigma-Aldrich, USA); 1% (vol/vol) B27 (#17504044, Invitrogen, Life Technologies, USA) and 6.2 g/l D-glucose. Slices were incubated in a 5% carbon dioxide (CO2) humidified incubator at between 35 - 37°C. Recordings were made after 6-14 days in culture. Previous studies have shown that after 7 days in culture (equivalent to postnatal day 14) GABAergic signaling and E_GABA_ has sufficiently developed to a level resembling mature nervous tissue (Streit et al., 1989; Wright et al., 2017). For the impermeant anion experiment, the mouse organotypic brain slices were injected one day after culture with adeno-associated vector serotype 1 (AAV1) containing a double-floxed sequence for channelrhodopsin (ChR2) linked to a yellow fluorescent protein (YFP) tag driven by the elongation factor 1 promoter (UNC Vector Core, USA). The vector was diffusely injected into slices using a custom-built Openspritzer pressurized ejection device (Forman et al., 2017). Slices were left for 6 days in culture to allow for robust expression of ChR2-YFP in GAD2+ interneurons.

For recordings, slices were transferred to a submerged chamber which was perfused with artificial cerebro-spinal fluid (aCSF) bubbled with carbogen gas (95% oxgen:5% carbon dioxide). aCSF was composed of (in mM): NaCl (120); KCl (3); Mg_Cl_2 (2); CaCl2 (2); NaH2PO4 (1.2); NaHCO3 (23); D-Glucose (11) with pH adjusted to be between 7.35-7.40 using 0.1 mM NaOH. Neurons were visualized using a 20X or 60x water-immersion objective (Olympus, Japan) on a BX51WI upright microscope (Olympus, Japan). Widefield images were obtained using a Mightex CCE-B013-U CCD camera. For the optogenetic experiments, we used epifluorescence microscopy to select slices that exhibited strong ChR2-YFP expression throughout the hippocampal region. Micropipettes were prepared from borosilicate glass capillaries with an outer diameter of 1.20 mm and inner diameter of 0.69 mm (Warner Instruments, USA), using a horizontal puller (Sutter, USA). Recordings were made using Axopatch 1D and Axopatch 200B amplifiers and acquired using pClamp9 (Molecular Devices) or WinWCP software (University of Strathclyde, UK).

For the CCC blockade experiment gramicidin perforated patch recordings (Kyrozis and Reichling, 1995) were performed using glass pipettes containing (in mM): 135 KCl, 4 Na2ATP, 0.3 Na3GTP, 2 Mg_Cl_2, 10 HEPES and 80 μg/ml gramicidin (Calbiochem; pH 7.38; osmolarity, 290 mosmol/l). After obtaining a cell-attached patch, the gramicidin perforation process was evaluated by continuously monitoring the decrease in access resistance. Recordings were started once the access resistance had stabilized between 20-80 MΩ, which usually occurred 20-40 min following gigaseal formation. Rupture of the gramicidin patch, referred to as a break through, induces a large influx of the high Cl^−^ internal solution into the cell. This causes a significant and permanent increase in the E_GABA_ at which time recordings would be discarded. GABA_A_R activation was achieved by pressure application of muscimol (10 μM, Tocris, UK), a selective GABA_A_R agonist, via a picrospritzer. To calculate E_GABA_, GABA_A_R currents were elicited at different command voltages. These were a series of 10 mV steps above and below −60 mV. Reported membrane potentials were corrected for the voltage drop across the series resistance for each neuron. Holding current (reflecting membrane current) and total current (reflecting membrane current plus the GABA_A_R-evoked current) were plotted against the corrected holding potential to generate a current-voltage (I-V) curve. Using this graph, the E_GABA_ was defined as the potential where the total current equals the holding current. Some of the data used for calculating E_GABA_ values in Figure 3E was also used in a previous study (Wright et al., 2017). V_m_ was defined as the x-intercept of the holding current, and the driving force as the difference between the two. To block CCC function, the KCC2 blocker furosemide (1mM, Sigma, USA) was applied.

For the impermeant anion experiments, whole-cell recordings were utilized as electroporation consistently ruptured the gramicidin patches. Pipettes were filled with internal solution composed of (in mM): Kgluconate (120); KCl (10); Na2ATP (4); NaGTP (0.3); Na2-phosphocreatinine (10) and HEPES (10). To test the effect of introducing impermeant anions, we electroporated the anionic 10 000 MW Dextran Alexa-Flour 488 (Thermo Fisher, USA) using a separate pipette positioned near the soma of the patched cell. This molecule is a hydrophilic polysaccharide, which is both membrane impermeant and highly negatively charged. Pipettes were filled with a 5% dextran solution in phosphate buffered saline and voltage pulses (5-10 of 20 ms duration, 0.5-1 V) were applied using a stimulus isolator. Successful electroporation of the anionic dextran was confirmed visually by observing the cell fill with the fluorescent dye. Electroporation also resulted in immediate membrane depolarization, which recovered over a period of 1-5 mins. E_GABA_, V_m_ and driving force were calculated during voltage steps in voltage clamp mode (as above) with GABA_A_Rs activated via endogenous synaptic release of GABA using photo-activation of GAD2+ interneurons expressing ChR2-YFP with 100 ms pulses of blue light using a 470 nm LED (Thorlabs), in the presence of 5 μM CGP-35348 (Torcis Bioscience, UK) to block GABA_B_R activation. E_GABA_, V_m_ and driving force were calculated before and atleast 5 mins following electroporation to allow for stabilization of V_m_, E_GABA_ and driving force. The image of the dextran filled cell was acquired using a confocal microscope (LSM510 Meta NLO, Car Zeiss, Jena, Germany). Analysis was performed using custom written scripts in the MATLAB environment (MathWorks). Numeric results are presented as mean +/− SEM.

## Acknowledgements

We would like to thank Juha Voipio for critically reading the manuscript and providing helpful comments and suggestions.

## References

Ambros-Ingerson J, Holmes WR (2005) Analysis and comparison of morphological reconstructions of hippocampal field CA1 pyramidal cells. Hippocampus 15:302–315.

Andrew RD, Labron MW, Boehnke SE, Carnduff L, Kirov SA (2007) Physiological evidence that pyramidal neurons lack functional water channels. Cereb Cortex 17:787–802.

Armstrong CM (2003) The Na/K pump, Cl ion, and osmotic stabilization of cells. Proc Natl Acad Sci U S A 100:6257–6262.

Baker PF, Willis JS (1972) Inhibition of the sodium pump in squid giant axons by cardiac glycosides: dependence on extracellular ions and metabolism. J Physiol 224:463–475.

Ben-Ari Y (2002) Excitatory actions of GABA during development: the nature of the nurture. Nat Rev Neurosci 3:728–739.

Berglund K, Wen L, Dunbar RL, Feng G, Augustine GJ (2016) Optogenetic Visualization of Presynaptic Tonic Inhibition of Cerebellar Parallel Fibers. J Neurosci 36:5709–5723.

Burton RF (1983) The composition of animal cells: solutes contributing to osmotic pressure and charge balance. Comp Biochem Physiol B 76:663–671.

Campbell SL, Robel S, Cuddapah Va., Robert S, Buckingham SC, Kahle KT, Sontheimer H (2015) GABAergic disinhibition and impaired KCC2 cotransporter activity underlie tumor-associated epilepsy. Glia 63:23–36.

Charron FM, Blanchard MG, Lapointe JY (2006) Intracellular hypertonicity is responsible for water flux associated with Na+/glucose cotransport. Biophys J 90:3546–3554.

Coull JAM, Boudreau D, Bachand K, Prescott SA, Nault F, Sík A, De Koninck P, De Koninck Y (2003) Trans-synaptic shift in anion gradient in spinal lamina I neurons as a mechanism of neuropathic pain. Nature 424:938–942.

Dai J, Sheetz MP, Wan X, Morris CE (1998) Membrane Tension in Swelling and Shrinking Molluscan Neurons. J Neurosci 18:6681–6692.

Diarra A, Sheldon C, Church J (2001) In situ calibration and [H+] sensitivity of the fluorescent Na+ indicator SBFI. Am J Physiol Cell Physiol 280:C1623–C1633.

Dierkes PW, Wüsten HJ, Klees G, Müller A, Hochstrate P (2006) Ionic mechanism of ouabain-induced swelling of leech Retzius neurons. Pflugers Arch Eur J Physiol 452:25–35.

Dijkstra K, Hofmeijer J, van Gils S, van Putten MJAM (2016) A Biophysical Model for Cytotoxic Cell Swelling. J Neurosci in press:11881–11890.

Doyon N, Prescott S a, Castonguay A, Godin AG, Kröger H, De Koninck Y (2011) Efficacy of synaptic inhibition depends on multiple, dynamically interacting mechanisms implicated in chloride homeostasis. PLoS Comput Biol 7:e1002149.

Doyon N, Vinay L, Prescott SA, De Koninck Y (2016) Chloride Regulation: A Dynamic Equilibrium Crucial for Synaptic Inhibition. Neuron 89:1157–1172.

Farrant M, Kaila K (2007) The cellular, molecular and ionic basis of GABAA receptor signalling. Prog Brain Res 160:59–87.

Ferrini F, Trang T, Mattioli TM, Laffray S, Del’Guidice T, Lorenzo L, Castonguay A, Doyon N, Zhang W, Godin AG, Mohr D, Beggs S, Vandal K, Beaulieu J, Cahill CM, Salter MW, De Koninck Y (2013) Morphine hyperalgesia gated through microglia-mediated disruption of neuronal Cl− homeostasis. Nat Neurosci 16:183–192.

Fettiplace R, Haydon DA (1980) Water permeability of lipid membranes. Physiol Rev 60:510–550.

Földy C, Lee S-H, Morgan RJ, Soltesz I (2010) Regulation of fast-spiking basket cell synapses by the chloride channel ClC-2. Nat Neurosci 13:1047–1049.

Forman CJ, Tomes H, Mbobo B, Burman RJ, Jacobs M, Baden T, Raimondo J V (2017) Openspritzer: an open hardware pressure ejection system for reliably delivering picolitre volumes. Sci Rep 7:2188.

Fraser JA, Huang CL-H (2004) A quantitative analysis of cell volume and resting potential determination and regulation in excitable cells. J Physiol 559:459–478.

Gagnon MP, Bissonnette P, Deslandes LM, Wallendorff B, Lapointe JY (2004) Glucose Accumulation Can Account for the Initial Water Flux Triggered by Na+/Glucose Cotransport. Biophys J 86:125–133.

Glykys J, Dzhala V, Egawa K, Balena T (2014) Local Impermeant Anions Establish the Neuronal Chloride Concentration. Science (80-):670–676.

Hamada K, Matsuura H, Sanada M, Toyoda F, Omatsu-Kanbe M, Kashiwagi A, Yasuda H (2003) Properties of the Na ^+^/K^+^ pump current in small neurons from adult rat dorsal root ganglia. Br J Pharmacol 138:1517–1527.

Hernández J a, Cristina E (1998) Modeling cell volume regulation in nonexcitable cells: the roles of the Na+ pump and of cotransport systems. Am J Physiol - Cell Physiol 275:C1067–C1080.

Hill T (1956) A. Fundamental studies. On the theory of the Donnan membrane equilibrium. Discuss Faraday Soc.

Hille B (2001) Ion Channels of Excitable Membranes. Sunderl Massachusetts USA.

Huberfeld G, Wittner L, Clemenceau S, Baulac M, Kaila K, Miles R, Rivera C (2007) Perturbed chloride homeostasis and GABAergic signaling in human temporal lobe epilepsy. J Neurosci 27:9866–9873.

Hyde TM, Lipska BK, Ali T, Mathew S V, Law AJ, Metitiri OE, Straub RE, Ye T, Colantuoni C, Herman MM, Bigelow LB, Weinberger DR, Kleinman JE (2011) Expression of GABA signaling molecules KCC2, NKCC1, and GAD1 in cortical development and schizophrenia. J Neurosci 31:11088–11095.

Jedlicka P, Deller T, Gutkin BS, Backus KH (2011) Activity-dependent intracellular chloride accumulation and diffusion controls GABA(A) receptor-mediated synaptic transmission. Hippocampus 21:885–898.

Jiang C, Haddad GG (1991) Effect of anoxia on intracellular and extracellular potassium activity in hypoglossal neurons in vitro. J Neurophysiol 66:103–111.

Kager H, Wadman WJ, Somjen GG (2000) Simulated seizures and spreading depression in a neuron model incorporating interstitial space and ion concentrations. J Neurophysiol 84:495–512.

Kaila K, Price TJ, Payne J a., Puskarjov M, Voipio J (2014) Cation-chloride cotransporters in neuronal development, plasticity and disease. Nat Rev Neurosci 15:637–654.

Kay AR (2017) How Cells Can Control Their Size by Pumping Ions. Front Cell Dev Biol 5:1–14.

Keener J, Sneyd J (1998) Mathematical Physiology, Interdisciplinary Applied Mathematics 8.

Klein PM, Lu A, Harper ME, McKown HM, Morgan J, Beenhakker MP (2017) Tenuous inhibitory GABAergic signaling in the reticular thalamus. J Neurosci:1345–17.

Krishnan GP, Bazhenov M (2011) Ionic dynamics mediate spontaneous termination of seizures and postictal depression state. J Neurosci 31:8870–8882.

Kuner T, Augustine GJ (2000) A Genetically Encoded Ratiometric Indicator for Chloride:: Capturing Chloride Transients in Cultured Hippocampal Neurons. Neuron 27:447–459.

Kyrozis A, Reichling DB (1995) Perforated-patch recording with gramicidin avoids artifactual changes in intracellular chloride concentration. J Neurosci Meth 57:27–35.

Lagostena L, Rosato-Siri M, D’Onofrio M, Brandi R, Arisi I, Capsoni S, Franzot J, Cattaneo A, Cherubini E (2010) In the Adult Hippocampus, Chronic Nerve Growth Factor Deprivation Shifts GABAergic Signaling from the Hyperpolarizing to the Depolarizing Direction. J Neurosci 30:885–893.

Lee HHC, Deeb TZ, Walker JA, Davies PA, Moss SJ (2011) NMDA receptor activity downregulates KCC2 resulting in depolarizing GABAA receptor–mediated currents. Nat Neurosci 14:736–743.

Lewin N, Aksay E, Clancy CE (2012) Computational Modeling Reveals Dendritic Origins of GABA A - Mediated Excitation in CA1 Pyramidal Neurons. 7.

Liang D, Bhatta S, Volodymyr G, Simard JM (2007) Cytotoxic edema : mechanisms of pathological cell swelling. Neurosurg Focus 22:1–9.

Lodish H, Berk A, Zipursky SL, Matsudaira P, Baltimore D, Darnell J (2009) Intracellular Ion Environment and Membrane Electric Potential. In: Molecular Cell Biology, 4th ed. New York.

MacAulay N, Gether U, Klærke DA, Zeuthen T (2002) Passive water and urea permeability of a human Na+-glutamate cotransporter expressed in Xenopus oocytes. J Physiol 542:817–828.

MacKenzie G, Maguire J (2015) Chronic stress shifts the GABA reversal potential in the hippocampus and increases seizure susceptibility. Epilepsy Res 109:13–27.

Mahadevan V, Dargaei Z, Ivakine EA, Hartmann A-M, Ng D, Chevrier J, Ormond J, Nothwang HG, McInnes RR, Woodin MA (2015) Neto2-null mice have impaired GABAergic inhibition and are susceptible to seizures. Front Cell Neurosci 9.

Misgeld U, Deisz R a, Dodt HU, Lux HD (1986) The role of chloride transport in postsynaptic inhibition of hippocampal neurons. Science 232:1413–1415.

Mohapatra N, Tønnesen J, Vlachos A, Kuner T, Deller T, Nägerl UV, Santamaria F, Jedlicka P (2016) Spines slow down dendritic chloride diffusion and affect short-term ionic plasticity of GABAergic inhibition. Sci Rep 6:23196.

Nichol JA, Hutter F (1996) Tensile strength and dilatational elasticity of giant sarcolemmal vesicles shed from rabbit muscle. J Physiol 493:187–198.

Price TJ, Cervero F, Gold MS, Hammond DL, Prescott SA (2009) Chloride regulation in the pain pathway. Brain Res Rev 60:149–170.

Qian N, Sejnowski TJ (1989) An Electro-Diffusion Model for Computing Membrane Potentials and Ionic Concentrations in Branching Dendrites, Spines and Axons. Biol Cybern 62:1–15.

Qian N, Sejnowski TJ (1990) When is an inhibitory synapse effective? Proc Natl Acad Sci USA 87:8145–8149.

Raimondo J V., Burman RJ, Katz A a., Akerman CJ (2015) Ion dynamics during seizures. Front Cell Neurosci 9:1–14.

Raimondo J V, Kay L, Ellender TJ, Akerman CJ (2012) Optogenetic silencing strategies differ in their effects on inhibitory synaptic transmission. Nat Neurosci 15:1102–1104.

Raimondo J V, Richards BA, Woodin MA (2017) Neuronal chloride and excitability — the big impact of small changes. Curr Opin Neurobiol 43:35–42.

Rivera C, Voipio J, Thomas-Crusells J, Li H, Emri Z, Sipilä S, Payne JA, Minichiello L, Saarma M, Kaila K (2004) Mechanism of activity-dependent downregulation of the neuron-specific K-Cl cotransporter KCC2. J Neurosci 24:4683–4691.

Rungta RL, Choi HB, Tyson JR, Malik A, Dissing-Olesen L, Lin PJC, Cain SM, Cullis PR, Snutch TP, Macvicar BA (2015) The cellular mechanisms of neuronal swelling underlying cytotoxic edema. Cell 161:610–621.

Sachs F, Sivaselvan M V. (2015) Cell volume control in three dimensions: Water movement without solute movement. J Gen Physiol 145:373–380.

Savtchenko LP, Poo MM, Rusakov DA (2017) Electrodiffusion phenomena in neuroscience: a neglected companion. Nat Publ Gr 18:598–612.

Sperelakis N (2012) Chapter 10 – Gibbs–Donnan Equilibrium Potentials. In: Cell Physiology Source Book, pp 147–151.

Staley KJ, Proctor WR (1999) Modulation of mammalian dendritic GABAA receptor function by the kinetics of Cl- and HCO3- transport. J Physiol 519:693.

Stoppini L, Buchs PA, Muller D (1991) A simple method for organotypic cultures of nervous tissue. J Neurosci Methods 37:173–182.

Streit P, Thompson SM, Gahwiler BH (1989) Anatomical and Physiological Properties of GABAergic Neurotransmission in Organotypic Slice Cultures of Rat Hippocampus. Eur J Neurosci 1:603–615.

Tang D, Qian A, Song D, Ben Q, Yao W, Sun J, Li W, Xu T, Yuan Y (2015) Role of the potassium chloride cotransporter isoform 2-mediated spinal chloride homeostasis in a rat model of visceral hypersensitivity. Am J Physiol Gastrointest Liver Physiol 308:G767–78.

Taniguchi H, He M, Wu P, Kim S, Paik R, Sugino K, Kvitsiani D, Kvitsani D, Fu Y, Lu J, Lin Y, Miyoshi G, Shima Y, Fishell G, Nelson SB, Huang ZJ (2011) A resource of Cre driver lines for genetic targeting of GABAergic neurons in cerebral cortex. Neuron 71:995–1013.

Thompson SM, Gähwiler BH (1989) Activity-dependent disinhibition. II. Effects of extracellular potassium, furosemide, and membrane potential on E_Cl_- in hippocampal CA3 neurons. J Neurophysiol 61:512–523.

Tosteson DC, Hoffman JF (1960) Regulation of cell volume by active cation transport in high and low potassium sheep red cells. J Gen Physiol 44:169–194.

Tyzio R, Minlebaev M, Rheims S, Ivanov A, Jorquera I, Holmes GL, Zilberter Y, Ben-Ari Y, Khazipov R (2008) Postnatal changes in somatic gamma-aminobutyric acid signalling in the rat hippocampus. Eur J Neurosci 27:2515–2528.

Tyzio R, Nardou R, Ferrari DC, Tsintsadze T, Shahrokhi A, Eftekhari S, Khalilov I, Tsintsadze V, Brouchoud C, Chazal G, Lemonnier E, Lozovaya N, Burnashev N, Ben-Ari Y (2014) Oxytocin- mediated GABA inhibition during delivery attenuates autism pathogenesis in rodent offspring. Science (80-) 343:675–679.

Voipio J, Boron WF, Jones SW, Hopfer U, Payne Ja, Kaila K (2014) Comment on “Local impermeant anions establish the neuronal chloride concentration”. Science 345:1130.

Wright R, Newey SE, Ilie A, Wefelmeyer W, Raimondo J V., Ginham R, Jeffrey Mcllhinney RA, Akerman CJ (2017) Neuronal chloride regulation via KCC2 is modulated through a GABAB receptor protein complex. J Neurosci 37:5447–5462.

Xiao A, Wei L, Xia S, Rothman S, Yu S (2002) Ionic mechanisms of ouabain-induced concurrent apoptosis and necrosis in individual cultured cortical neurons. J Neurosci 15:4.

Yeh BK, Lazzara R (1973) Reversal of Ouabain-lnduced Electrophysiological Effects by Potassium Canrenoate in Canine Purkinje Fibers. Circ Res 32:501–508.

Zeuthen T (1994) Cotransport of K+, Cl- and H2O by membrane proteins from choroid plexus epithelium of Necturus maculosus. J Physiol 478 (Pt 2:203–219.

